# Hepatic CD8^+^TOX^+^ T-cells are a hallmark of autoimmune hepatitis

**DOI:** 10.64898/2026.07.06.734562

**Authors:** Marc S. Sherman, Daniel M. Schafer, Molly F. Thomas, Sonya W. Katzen, Genevieve Boland, Angela R. Shih, Georg M. Lauer, Alexandra-Chloe Villani, Wolfram Goessling

## Abstract

Autoimmune hepatitis (AIH) is a chronic progressive liver disease that despite suggestive serum autoantibodies or plasma cell enrichment, remains functionally a diagnosis of exclusion. Whether the broader cellular composition of the liver might enable improved specificity of diagnosis has not been systematically tested. We prospectively recruited patients undergoing a clinically-indicated liver biopsy for suspected AIH and performed single-nucleus RNA sequencing (snRNA-seq) on biopsy tissue to map the cellular landscape of AIH and its diagnostic mimics. Unsupervised clustering on cell-type abundances alone largely separated AIH from non-AIH samples. Among individual populations, a subset of CD8⁺ T-cells marked by high TOX and PD1 expression was the most discriminating feature: its enrichment perfectly distinguished AIH by both snRNA-seq and *in situ* density (AUC = 1.00), outperforming plasma cell abundance (AUC = 0.83). CD8⁺TOX⁺ T-cell enrichment may therefore be the histologic lesion that marks the diagnosis of AIH.

## Introduction

Autoimmune hepatitis (AIH) is a chronic, immune-mediated liver disease characterized by interface hepatitis^1,2^, circulating autoantibody production^3^, and elevated serum transaminases^4,5^. Despite clinical consensus guidelines and scoring systems that incorporate demographic, serologic, biochemical, and histologic features^6,7^, diagnosis remains imprecise^8–10^. Autoantibodies, although central to diagnostic scoring, lack either sensitivity (anti-LKM1, anti-SLA) or specificity (anti-smooth muscle antibody [ASMA], anti-nuclear antibody [ANA]), and are frequently present in patients with metabolic liver disease and other autoimmune conditions^8–12^. Consequently, patients undergo an extended evaluation including a liver biopsy^13,14^ that delays diagnosis and treatment initiation^15^. Even then, some patients may be misclassified, with misclassification common enough that adequate response to treatment is a codified decision point in the revised original score for AIH^6^.

Histopathology is requisite for AIH diagnosis^13^ but it is also limited by subjective interpretation and incomplete specificity^6,16^. The presence of a lymphoplasmacytic infiltrate spilling across the portal hepatic limiting plate into the lobule is a well-recognized feature of AIH^6,17–20^, yet lymphocytes and plasma cell-rich infiltrates can be observed in drug-induced liver injury (DILI)^19,20^, viral hepatitis^17,20^, and steatohepatitis^21^. Interface hepatitis, historically considered a defining histopathologic feature of AIH, was recently reported to be present in 33% of cases with simple steatosis or early metabolic dysfunction-associated steatohepatitis (MASH), and in 100% of patients with MASH and advanced fibrosis^21^. Similarly, primary biliary cholangitis (PBC) may manifest with an interface hepatitis^22,23^, which when pronounced suggests AIH-PBC overlap syndrome^24–26^, an entity that may simply represent a particularly inflammatory form of PBC^25,26^. Other immune features like hepatic rosettes and emperipolesis are also now recognized to be hallmarks of hepatocyte injury more so than specific histologic markers of AIH^16^. As a result, histologic distinction between AIH and other inflammatory liver conditions remains challenging.

Beyond morphologic description, our understanding of the immunologic architecture in the liver of patients with AIH is patchy. Cell types reported to be enriched in AIH include CD3^+^ T-cells^19,27,28^, CD4^+^ T-cells^28–30^, CD8^+^ T-cells^19,28,29^, FOXP3^+^ cells (marking T_regs_)^28,29,31^, CD161^+^ cells (marking NK cells)^28^, mucosal-associated invariant T-cells (MAITs)^28,32^ and IgG^+^ plasma cells^29,33^, with skewing of IgG to IgM plasma cells in AIH versus PBC^33^. These studies span children and adults, and with controls largely limited to healthy tissue or specific disease comparators like PBC or MASH. These studies suggest specific cellular subtypes may mark the presence of AIH, and motivate the use of broad and unbiased methods to specify the immune landscape within a cohort that is diagnostically ambiguous at time of biopsy.

Tissue-based single-cell (scRNA-seq) and single-nucleus (snRNA-seq) RNA sequencing have identified the cellular and molecular signatures that specify other human autoimmune diseases like systemic lupus erythematosus^34^, multiple sclerosis^35,36^, and rheumatoid arthritis (RA)^37^. In RA, scRNA-seq from RA-inflamed synovial tissue enabled discrimination from osteoarthritis and further subclassification of synovia into cell-type abundance phenotypes that stratified treatment responsiveness to methotrexate or anti-TNF therapies^37^, demonstrating that despite human variation, clinically relevant insight may be gleaned from modestly-sized but well-pedigreed patient cohorts.

In this study, we prospectively recruited patients undergoing diagnostic evaluation for liver injury for whom the treating hepatologists suspected AIH. We used snRNA-seq to define the intrahepatic cellular landscape of AIH and compared it to MASH, PBC, overlap syndromes, resolving inflammation, and histologically normal liver. Unsupervised clustering algorithms trained on cell type abundances (CTAs) alone, without access to clinical, serologic, or demographic information, were largely sufficient to distinguish AIH from other liver disease states and identified CD8⁺TOX⁺ T-cells as a hallmark of AIH. In this discovery study, hepatic enrichment of CD8⁺TOX⁺ T-cells was diagnostic of AIH, while plasma cells were enriched in most patients with AIH but not disease defining. Our findings provide a data-driven, cellular definition of AIH, and suggest CD8⁺TOX⁺ T-cell enrichment may be the canonical histologic lesion of AIH.

## Results

### Cohort of treatment-naïve patients with AIH and disease controls

From September 2020 to June 2022, 30 unique patients were prospectively recruited and liver specimens were collected and successfully processed for snRNA-seq (Fig. 1a,b). 25 specimens originated from 24 patients with suspected AIH (1 re-biopsy). All biopsies were taken prior to therapy (“Diagnostic cohort”), and 6 specimens were from patients without underlying liver disease undergoing hepatic resection, primarily for colorectal metastasis (“Normal cohort”). Compared to the normal cohort, the diagnostic cohort was slightly younger and showed increased liver injury, as expected (Table 1). Patients tended to have ALT>AST, consistent with a cohort limited in alcohol use, and positive ASMA or ANA testing, consistent with patients suspected of having AIH. Diagnoses at time of biopsy (Table 1) highlighted metabolic dysfunction-associated steatotic liver disease (MASLD) or MASH as the major non-AIH diagnosis. Non-MASLD cases included AIH-PBC overlap, AMA-negative PBC, DILI and resolving liver injury.

**Table 1.**
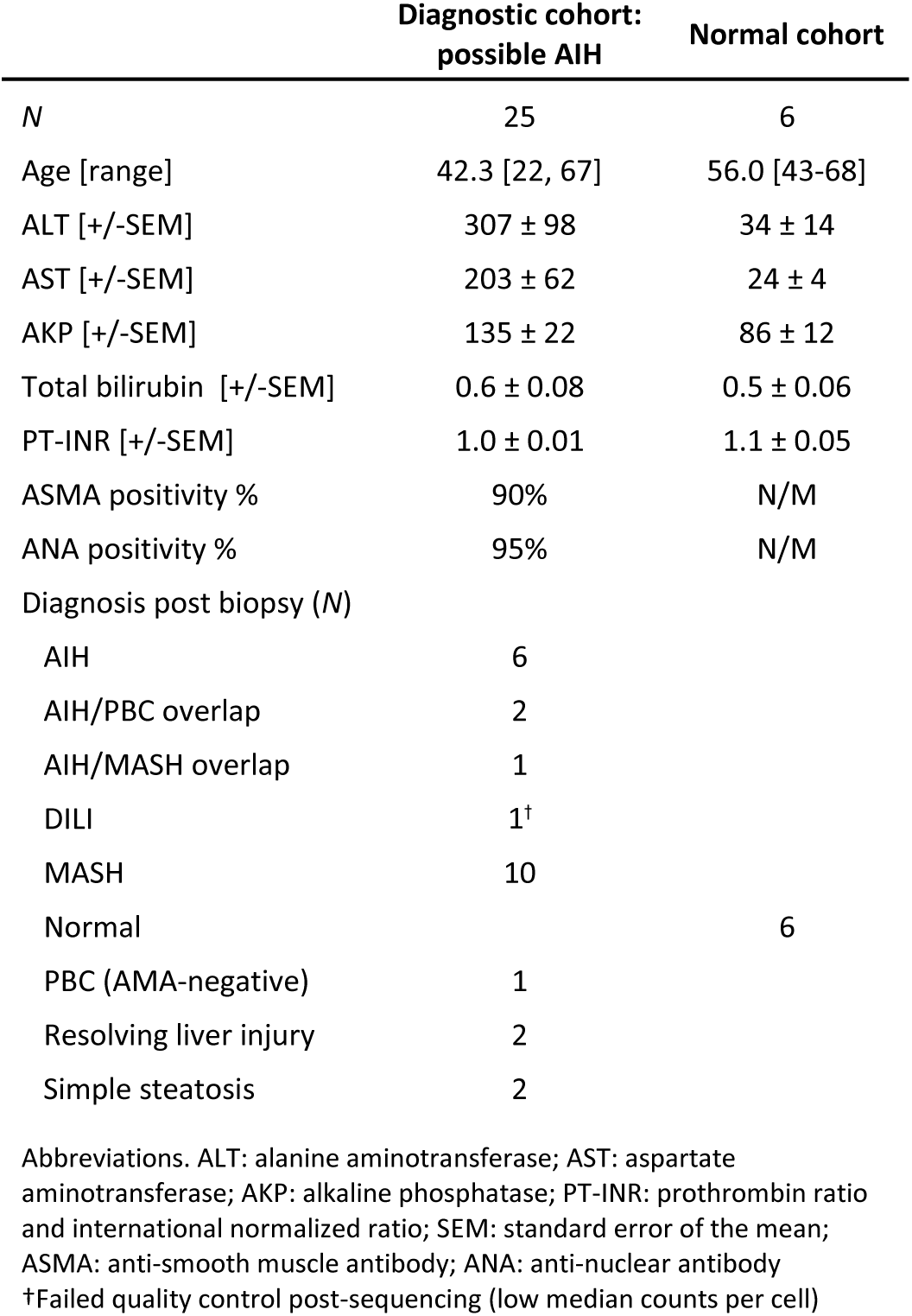

**Figure 1.**
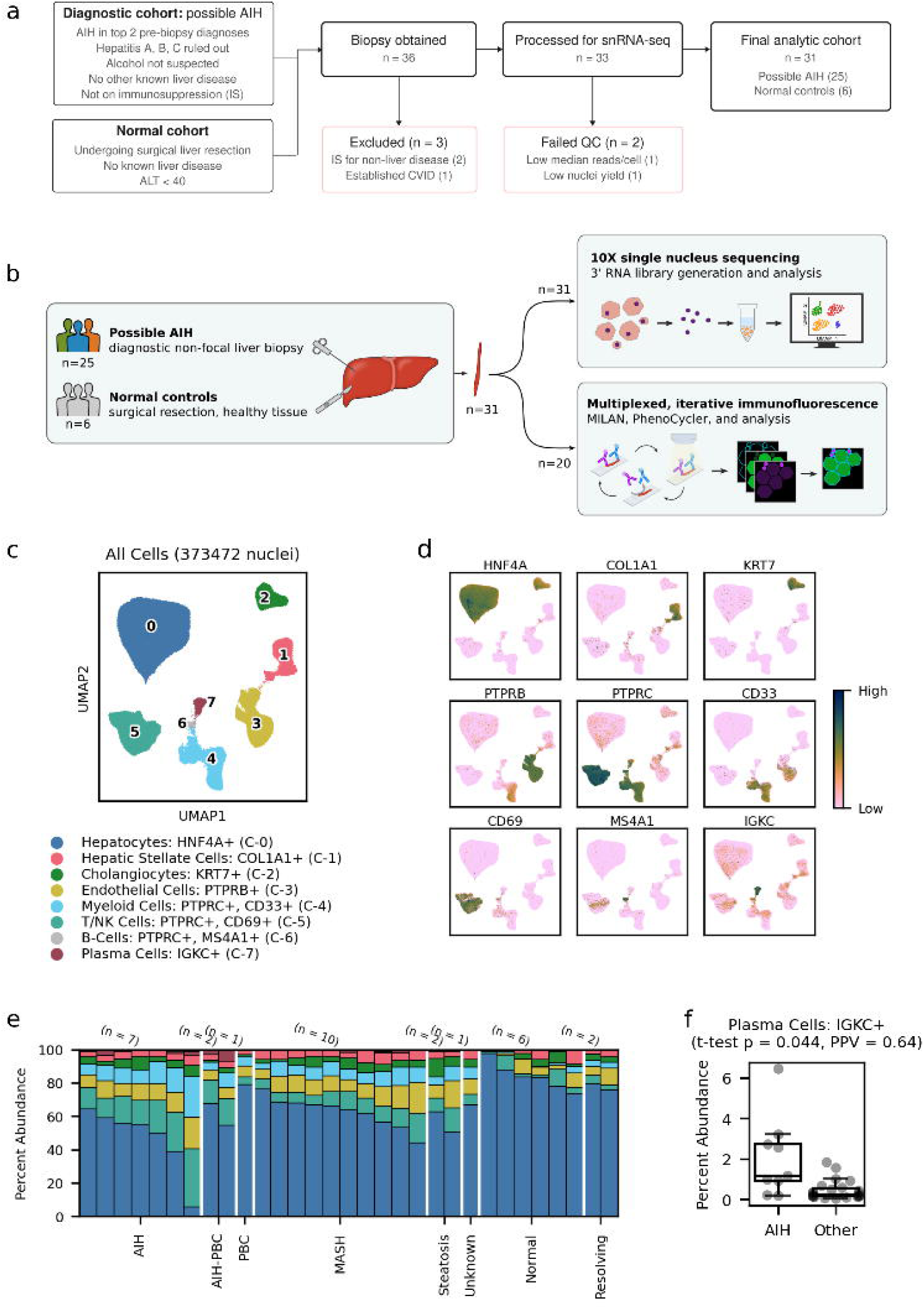
Prospective recruitment of patients with possible AIH study design, tissue processing, snRNA-seq, and elaboration of patient CTAs. (a) Prospective study design overview and attrition. (b) Tissue acquisition and sample processing for snRNA-seq (primary study) or multiplexed, iterative immunofluorescence (subset for internal validation). (c) All-nuclei UMAP, with clustering (Leiden) to reveal coarse cell types and our annotations. (d) UMAP expression of selected marker genes from pink (low expression) to green (high expression). (e) Coarse cell type fractional abundances for each sample, grouped by final diagnosis (x-axis). (f) Number of plasma cells divided by total cells per sample as a percent for non-AIH versus AIH, using inclusive AIH definition. Each dot is a sample; boxplot shows 25th, 50th and 75th quartiles, from bottom to top; whiskers show data range excluding outliers. Welch’s t-test reports a significant difference between AIH and non-AIH samples (p=0.044). Cutoff for positive predictive value (PPV) is determined by Youden’s J optimal cutoff.

Median clinical follow-up of the diagnostic cohort was 3.46 years (range [0.22, 4.05]), during which the diagnosis per the treating hepatologist changed for 2 patients (8%) (Table 2). One patient (S21) with comorbid Crohn’s disease on infliximab was presumed to have AIH-like infliximab-mediated DILI^38–40^ but her biopsy also met criteria for MASH. Her liver injury resolved with infliximab discontinuation and a coarse of budesonide, but relapsed after budesonide cessation, leading to a second biopsy (S50) which affirmed relapse of AIH features. She subsequently responded to budesonide induction and azathioprine maintenance, and, given complete biochemical resolution of liver injury despite weight gain was deemed to have AIH without clinically significant MASH. The second patient (S45) had Hashimoto’s thyroiditis and Sjogren’s disease, a BMI of 53, and severe chronic liver injury (ALT 591 IU/L pre-biopsy) and was found to have an equivocal liver biopsy showing a mild lymphoplasmacytic infiltrate without evidence of MASLD/MASH (no steatosis, ballooning, or sinusoidal fibrosis). Corticosteroid treatment led to worsening liver injury (peak ALT of 1283 IU/L), while corticosteroid cessation improved her liver injury to pre-biopsy baseline. She subsequently underwent previously planned bariatric surgery, after which her liver injury entirely ceased (ALT of 11 IU/L), suggesting a metabolic etiology despite not meeting histological criteria for MASLD.

Using the final, post-follow up diagnoses as ground truth, the index clinical characteristics of patients ultimately diagnosed with AIH or overlap syndromes did not substantially differ from non-AIH patients (Table 3). Among documented clinical demographic and laboratory values, only age (lower in non-AIH) and AST (higher in AIH) were significant (Table 3). Total IgG trended higher in AIH than non-AIH (1523 vs 1400 mg/dL, p=0.48), though this trend was almost entirely driven by high total IgG in the AIH-PBC overlap patients (2067.5 ± 185.5 mg/dL); and while ASMA antibody titers trended higher in AIH, ANA titers trended higher in non-AIH disease controls. Patient BMI was not discriminating, even considering AIH versus MASH.

Both the revised original AIH score and simplified scores were also calculated. The simplified AIH score^7^ was not significantly higher in AIH vs non-AIH (4.9 vs 4.5, p=0.50), and was actually lower in the AIH group when excluding the overlap cases (4.2 vs 4.5, p=0.52). The revised original AIH score^6^ was significantly higher in AIH diagnoses versus non-, regardless of precise comparison made (p<.005 all comparisons). However, the area under the curve (AUC) for the revised original score was 0.85 (Supp. Fig. 1), corresponding to a sensitivity of 1.0, a specificity of 0.68, and a positive predictive value of 0.64 (Youden’s J), in agreement with a robust prior independent validation assessment (sensitivity 1.0, specificity 0.73, positive predictive value 0.67^8^).

Collectively, this real-world, prospective study of patients with possible AIH confirmed by long-term follow-up highlights poor discrimination of clinical characteristics for AIH, and sensitive but relatively non-specific scoring for AIH. Post-biopsy, the major competing diagnosis was MASH. Although MASH was distinguishable from AIH histologically, two diagnoses yet changed after extensive follow up, both in whom MASH was the competing diagnosis. These data underscore the importance of identifying disease-defining features of AIH.

### snRNA-seq identifies major lineages and sub-lineages of the liver from fractional needle core biopsy

Liver biopsy specimens were sufficient input for snRNA-seq and enabled specification of all major lineages and sub-lineages of the liver (Fig. 1b). After quality control and batch-processing (Supp. Fig. 2), nuclei uniformly mapped to cell type and not patient sample (Supp. Fig. 2d). Recovered coarse cell types (Fig. 1c,d) included hepatocytes (*HNF4A*^+^, C-0), hepatic stellate cells (*COL1A1*^+^, C-1), cholangiocytes (*KRT7*^+^, C-2), endothelial cells (*PTPRB*^+^, C-3), and diverse *CD45*^+^ (*PTPRC*+, C-4, C-5, C-6) lineages corresponding to myeloid and lymphoid lineages, including B- (*CD20*^+^/*MS4A1*^+^, C-6) and plasma cells (*IGKC*^+^, C-7). Sample level abundances of coarse cell types showed marked changes between healthy or resolving injury and chronic disease states (Fig 1e). In particular, AIH and AIH-PBC overlap samples demonstrated expansion of lymphoid>myeloid lineages over normal and disease controls, though there was considerable variation. Plasma cells, considered a histologic hallmark of AIH^41–44^, were significantly increased but not discriminatory (t-test p=0.044; positive predictive value [PPV]=0.64); only 4 AIH samples showed fractional plasma cell enrichment higher than the highest individuals in the non-AIH category (Fig. 1f).

We resolved the lymphoid (T/NK), myeloid, and B/plasma compartments at high granularity, identifying 34 populations across these three superclusters. Complete annotation including cluster-defining markers and supporting evidence for every population is provided in the Supplementary Information (“Annotation of snRNA-seq cell types”; Supp. Fig. 3). Here we focus on the populations differentially abundant in AIH.

We identified 15 populations of T- and NK cells marked by distinct combinations in the expression of CD3, alpha-beta T-cell receptor (TCR) (*TRAC*, *TRBC1*, *TRBC2*), gamma-delta TCR (*TRDC*), CD4, and CD8 (Fig. 2a-c). Relevant for subsequent analyses were T-regulatory (T_reg_) cells (T-2), identified by their enrichment of *CD4*, *FOXP3*, *IKZF2*, *IL2RA* (CD25), and *CTLA4*^45,46^ and MAIT cells (T-7), identified by *SLC4A10*, *IL7R*, *IL23R*, and *KLRB1*^47,48^ (Fig. 2b,c). In addition, a prominent CD8+ T-cell population (T-9) was highlighted by the activation markers *LAG3*, *TOX*, and *PD1*^49,50^.

**Figure 2.**
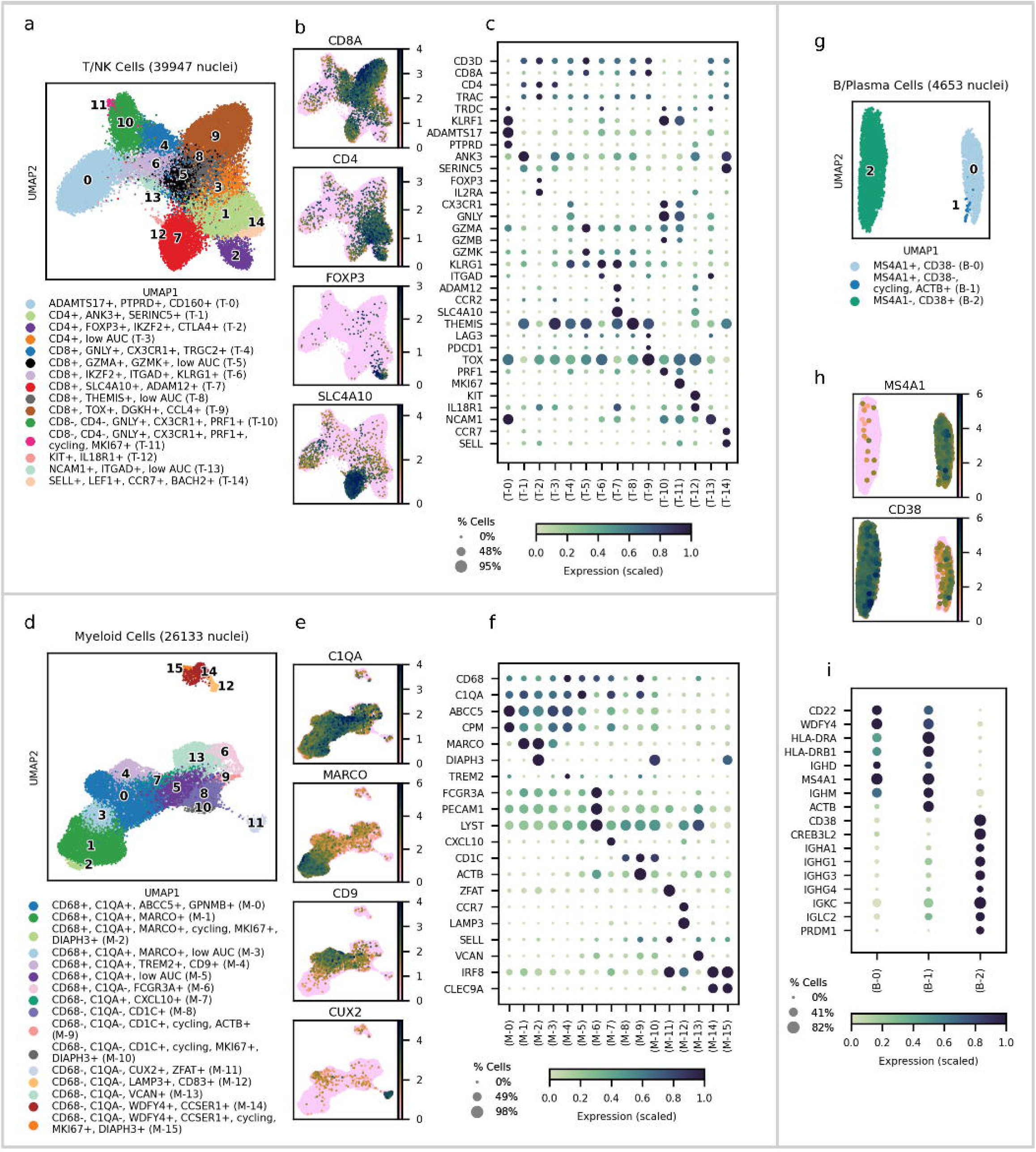
Coarse immune cell types (T/NK cells, Myeloid cells, and B-cells) were individually clustered to identify sub-cell types. UMAPs for lymphoid (a), myeloid (d), and B/plasma (g) lineages colored by fine-grained subtypes. The “low AUC” label indicates clusters that were not well-defined by marker gene analysis. Single-marker UMAP overlays for lymphoid (b), myeloid (e), and B/plasma (h). Marker gene dotplot for lymphoid (c), myeloid (f), and B/plasma (i) lineages. Gene expression goes from pink (low expression) to green (high expression). Gene expression is min-max normalized across samples and goes from green-blue (low expression) to dark blue (high expression).

Four major myeloid subtypes were identified: Kupffer cells (M-1, M-2), monocyte-derived macrophages (M-0,M-3, M-4, M-5), dendritic cells (DCs; M-7, M-8, M-9, M-10, M-11, M-12, M-14, M-15), and monocytes (M-6, M-13) (Fig. 2d). Among these, we highlight the DC populations and one monocyte-derived macrophage population. Conventional DC1 (cDC1) cells were distinguished by enriched expression of *IRF8*, *CLEC9A*, *WDFY4*, *CCSER1*, and *XCR1*, while cDC2 cells (M-8, M-9, M-10) were marked by *CD1C*, *CLEC10A* and *FLT3*^51,52^. Two other DC or DC-like populations (M-7, M-12) were identified and shared gene signatures. Cluster M-12 was *LAMP3*^+^ and colocalized with cDC1 (M-14), but was distinguished by expression of PD-L1/*CD274* and high *CCR7*, consistent with the recently described “mature DCs enriched in immunoregulatory molecules” (mregDCs)^53^, which may represent an activated cell state of cDC1 or cDC2^54–56^. Cluster M-7 was adjacent to M-4 and M-0 suggesting a macrophage lineage, but shared the canonical DC marker *LAMP3* (intermediate expression). M-7 was particularly high in T-cell chemoattractants *CXCL9*, *CXCL10*, and *CXCL11*, and shared features with a putative macrophage population (hM∅_9_) previously noted^56^. Among monocyte-derived macrophages, of note was a TREM2^+^CD9^+^ population (M-4) that shared characteristics with the recently described scar-associated macrophage (SAM) implicated in MASH^57,58^.

These results confirm capture of all major, and many minor parenchymal and non-parenchymal cell populations of human liver in health and disease, and are publicly available on an atlas/browser (https://liverquant.mgh.harvard.edu). Given this detailed immune taxonomy, we asked whether immune subpopulations were discriminatory for the diagnosis of AIH.

### Unsupervised clustering of patient hepatic cell type abundances distinguishes AIH from disease controls

CTAs, both at the coarse level (Fig. 1e; Supp. Fig. 4a) and within lineages (Fig. 3a; Supp. Fig. 4b-g), varied both between and within diagnostic categories. Although some patterns emerged by inspection, like expansion of lymphoid and myeloid populations in AIH versus controls, there was substantial within- and between-disease heterogeneity. We asked whether an unsupervised clustering approach of a combination of coarse and fine-mapped immune CTAs (Fig. 3a,b) could reveal underlying patterns in patient similarity. Crucially, unsupervised clustering on CTAs omits patient diagnostic labels and does not involve training.

**Figure 3.**
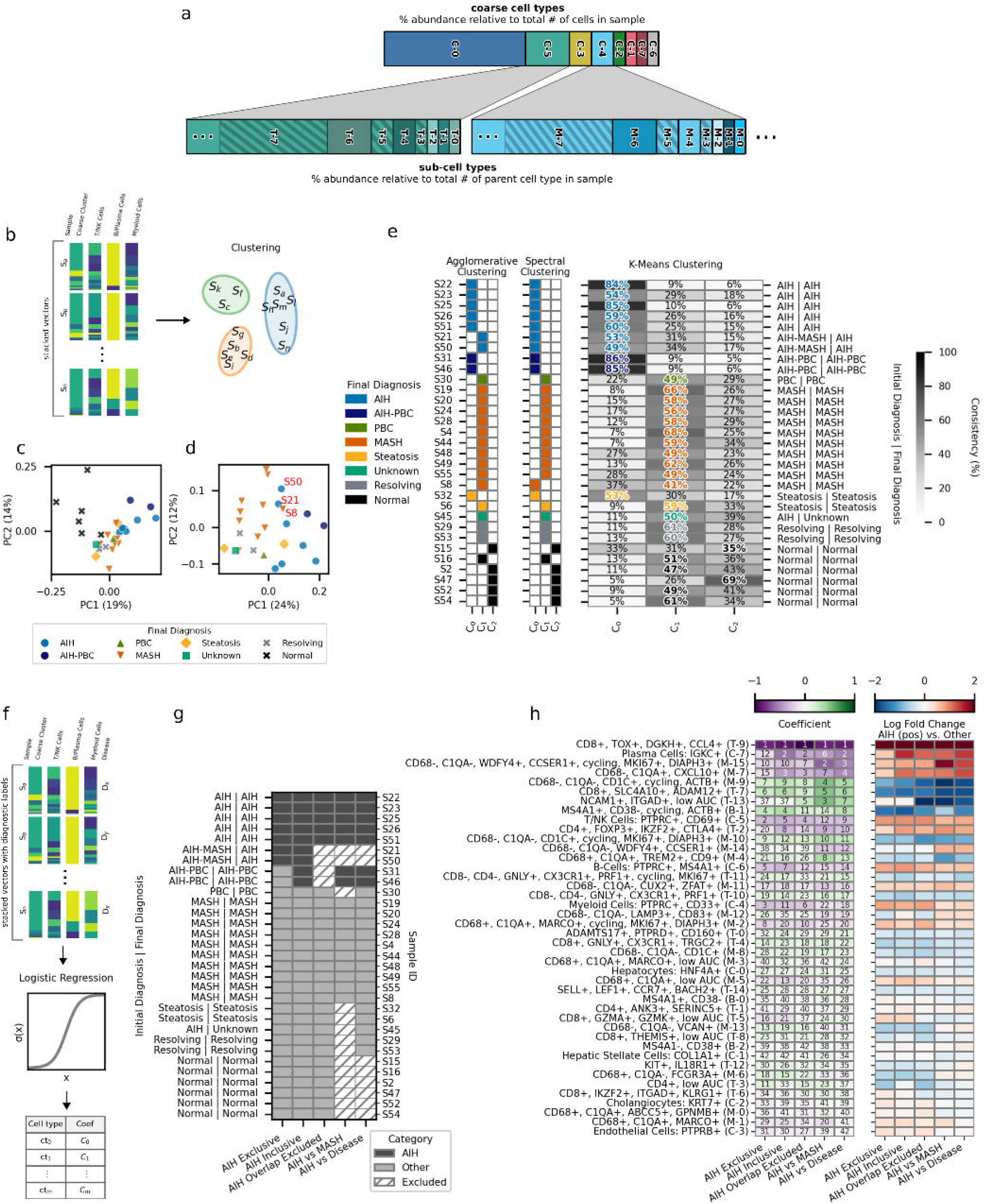
AIH is distinguished from non-AIH by naive clustering of coarse and immune-subtype abundances. (a) Anatomy of a CTA; a coarse CTA vector is composed of the fractional abundance of coarse cell type counts normalized by total cell count per patient. Similarly, a subtype CTA is a subtype cell count normalized by the total number of that coarse category (e.g., myeloid cells). (b) Overview of naive clustering: coarse and immune subtype CTAs are combined into a vector for each patient (or sample) S_x. Clustering then operates on a per-sample basis, without encoding patient diagnosis. (c) PCA of stacked CTAs separates normal (black) from diseased; (d) excluding normals, patients largely separate by AIH (right) versus non-AIH (left). (e) Naive clustering using agglomerative (left), spectral (middle), and k-means consistently reveals separation of AIH samples (C_0) from diseased non-AIH (C_1) and normal (C_2). Spectral and agglomerative clustering consistently reported the same clusters across different random seeds, therefore consistency is not reported. K-means percentages reflect cluster assignment across 5000 iterations for different random seeds. (f) Diagram demonstrating logistic regression analysis; stacked coarse and immune subtype CTA vectors coupled with binary diagnosis labels were used to train a logistic regression model. Model coefficients were retrieved for interpretation. (g) Binary disease labels are defined five ways in sensitivity analysis. The primary analysis was considered AIH versus disease omitting an AIH-MASH overlap (right-most column). (h) Coefficients from logistic regression according to case-control definition, ranked by the absolute value of coefficients from the model trained on AIH vs Disease defined labels.

To understand if such formal clustering might be fruitful, we first performed principal component analysis (PCA) on patient coarse and immune CTAs to visualize relationships among samples in two dimensions. The clearest separation of samples was between normal and diseased diagnoses (Fig. 3c). After removing normal samples, a separation between AIH (including AIH-PBC overlap) and other diagnoses emerged (Fig. 3d). The AMA-negative PBC patient localized to the interface of AIH and MASH clusters, and far from the AIH-PBC cases. As a consistency check, both biopsies of our index diagnosis AIH-MASH overlap patient, taken months apart (S21 initial biopsy, S50 second biopsy, both off-treatment) clustered together and fell near the intersection of the AIH and MASH clusters. These findings show reproducibility of CTAs produced by snRNA-seq and separation by disease state, thus motivating a more formal clustering analysis to capture subtle, multidimensional patterns.

We evaluated three clustering algorithms: spectral, agglomerative, and k-means (Fig. 3e). Performance with three clusters demonstrated consistent and biologically relevant results for spectral and agglomerative clustering (Supp. Fig. 5). All algorithms created an AIH-majority cluster that included AIH-PBC overlap patients and largely excluded patients with other diagnoses except steatosis sample S32 (C_0_). All algorithms also created a second non-AIH majority cluster (C_1_) comprised primarily of MASH patients. Spectral and agglomerative clustering created a third cluster containing most normal samples while excluding other diseased samples (C_2_). K-means failed to generate a convincing normal-majority group but still created a normal-exclusive cluster with S15 and S47. The patient exhibiting an AIH-MASH phenotype (S21, S50) on index biopsy as well as a MASH patient (S8) switched between C_0_ and C_1_ across spectral and agglomerative clustering. In k-means, these three samples (S21, S50, S8) also displayed similar inconsistency with higher rates of dual appearance in C_0_ and C_1_ compared to all other samples. The same samples also localized to the AIH-MASH boundary by PCA (Fig. 3d).

These naïve clustering approaches imply test characteristics with positive predictive values of 0.875 (agglomerative), 0.82 (spectral), and 0.91 (k-means), with modest (agglomerative) or no loss of sensitivity. Thus, while clustering was imperfect, it also derived solely from objective biopsy characteristics, and improved upon established diagnostic scoring criteria. Together, these data suggest that biopsy-derived fractional representation of coarse and fine cell immune types may improve the diagnosis of AIH and warranted further investigation.

### Cell types driving discrimination of AIH from disease controls

To understand the cell types driving co-clustering of AIH cases, we used coefficient values from a trained logistic regression model as a proxy for the predictive capacity of a cell type’s abundance (Fig. 3f). The logistic regression model was trained with all samples, using CTAs as input and binary diagnoses as labels. Multiple binary label sets were defined to evaluate the robustness of cell type contributors across differing definitions of case versus control (Fig. 3g). Coefficients favoring AIH (negative) or other (positive) were ranked by absolute magnitude (Fig. 3h). Remarkably, fractional enrichment of CD8^+^TOX^+^ T-cells (T-9) led the ranking as the most crucial feature defining the AIH irrespective of the precise case-control definition. Plasma cell enrichment, a hallmark of AIH, was ranked highly among the coefficients, but had a lower average log-fold change and was more inconsistent across varying label definitions compared to CD8^+^TOX^+^ T-cells.

This analysis revealed other patterns that define AIH versus non-AIH. AIH was favored by relative expansion of the DC-like *CXCL10*^HI^ (M-7) population, expansion of putative cDC1 (M-14), especially their cycling form (M-15), and contraction of putative cDC2 (M-9, M-10, M-8). Total T/NK cell abundances (C-5) ranked highly favoring AIH, but were not as specific as the TOX^+^CD8^+^ T-cell subset. Lastly, FOXP3^+^CD4^+^ T-cells, putatively T_regs_, were enriched in AIH as previously reported^31^. Interestingly, MAIT cell depletion was predictive of AIH (Supp. Fig. 6a,b), despite prior evidence of their enrichment in AIH^28^, suggesting that other T-cells might expand more robustly than MAIT cells. Finally, putative SAMs (M-4) were also depleted in AIH, and enriched particularly in MASH (Supp. Fig. 4g).

### AIH is defined by enrichment in CD8^+^TOX^+^ T-cells

Given the dominant influence of the CD8^+^TOX^+^ T-cell enrichment in driving clustering of AIH cases, we further evaluated this compartment and its relation to the AIH label. Among several ways one could construct the CTA vector, we had chosen coarse cell types appended to sub-cell types (Fig. 3a,b), which means the predictive power of CD8⁺TOX⁺ T-cells was relative to total T/NK fraction. Visualizing this measure directly, CD8^+^TOX^+^ T-cells were enriched in AIH, but not diagnostic (Supp. Fig. 6c), with two non-AIH patients having similar enrichment as the AIH patients.

Remarkably however, CD8^+^TOX^+^ T-cell abundance normalized to total cells (per sample) was perfectly discriminating for AIH (AUC=1.0) (Fig. 4a). Among other cell types enriched in AIH from the logistic regression, total fractional enrichment of presumptive T_regs_ (AUC=0.98), total T/NK cells (AUC=0.95), and total CD8^+^ T-cells (AUC=0.94) were also superior to fractional enrichment of plasma cells (AUC=0.83) (Fig. 4b-e).

**Figure 4.**
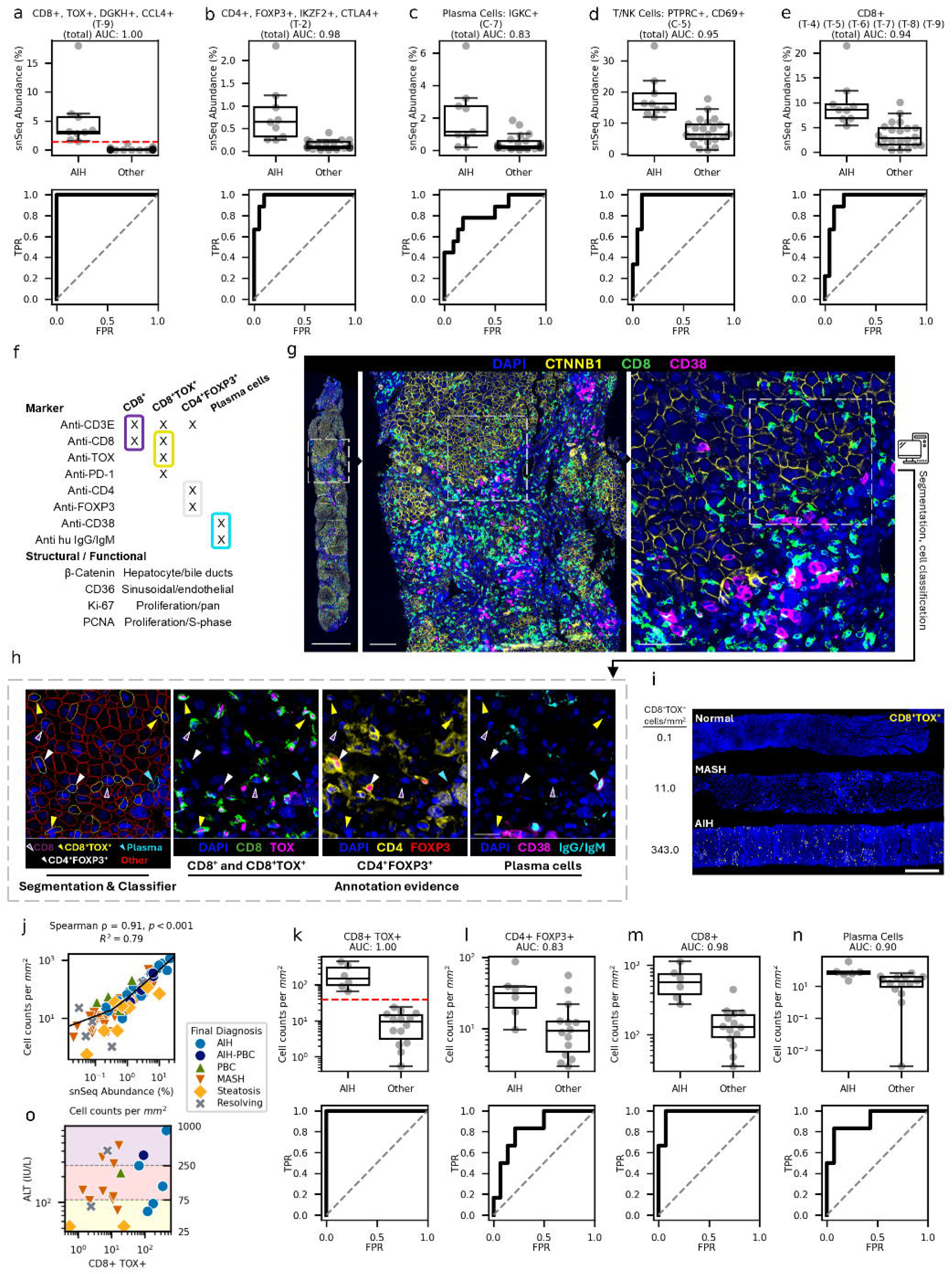
Enrichment of CD8^+^TOX^+^ T-cells (T-9) is diagnostic of human AIH. (a-e, top row) AIH versus non-AIH (other) fractional abundance of specific cell types divided by total cells (per sample), using the inclusive definition of AIH. Red dashed line (a) indicates an abundance threshold with perfect discrimination between AIH and non-AIH. (a-e, bottom row) Receiver operating characteristic curves for each cell type, with area under the curve (AUC) in the title indicating overall performance. Using a custom staining panel (f) liver biopsy specimens were imaged whole-slide at 20X magnification (g) using PhenoCycler (scalebars: (left) 1mm, (middle) 100μm, (right) 50μm). Cells were segmented and annotated (h) according to the boxed double positive markers in (f), with the arrows marking examples of CD8+ T-cells (purple, white outline), CD8^+^TOX^+^ T-cells (yellow), CD4^+^FOXP3^+^ T-cells (white), and plasma cells (cyan) (scalebar: 25μm). (i) Example of CD8^+^TOX^+^ T-cells (yellow) on DAPI background (blue) showing abundance differences between diagnoses, where the sample selected was the patient with median CD8^+^TOX^+^ T-cell abundance for Normal, MASH and AIH (scalebar: 800μm). (j) snSeq relative abundance (x-axis) versus area density abundance (y-axis), where relative abundance was snSeq relative frequency of each cell type divided by total cells per sample, while area density abundance was PhenoCycler annotated cells per unit tissue area. (k-n, top row) AIH versus non-AIH (other) area density abundance of specific cell types (per sample), using the inclusive definition of AIH. (k-n, bottom row) Receiver operating characteristic for associated plots, with AUC in the titles. (o) Spatial area density of CD8^+^TOX^+^ T-cells (x-axis) versus ALT from time of biopsy (y-axis), colored by final diagnosis.

One advantage of snRNA-seq (over scRNA-seq) is that the overall cell type abundance is thought to hew closely to the *in situ* abundances^59^. We therefore hypothesized that (1) snRNA-seq fractional representation predicts unit area measurement *in situ* (hereafter denoted as area density), and that (2) CD8^+^TOX^+^ T-cells area density enrichment is diagnostically discriminating for AIH.

To operationalize measurement of specific cell types we prototyped an antibody panel for identifying CD8^+^TOX^+^ T-cells and other discriminating cell types *in situ* (Fig. 4f) (see Methods). The panel was applied to patients from the snRNA-seq cohort using the Akoya PhenoCycler platform to reveal the spatial distributions and area densities of relevant immune populations (Fig. 4f-i). Cells were segmented using InstansSeg^60^ (Fig. 4h). Two spatially colocalized markers were required to denote the presence of specific cell types (see Fig. 4f,h). A multi-label classifier was trained to identify CD8^+^ T-cells, CD8^+^TOX^+^ T-cells, CD4^+^FOXP3^+^ cells, and plasma cells (see Methods).

Area density measurements of putatively AIH-enriched cell types (Fig. 4j) revealed remarkable concordance with our snRNA-seq abundance (Spearman ρ=0.91, p<.001). First, numerical enrichment of CD8^+^TOX^+^ T-cells perfectly distinguished AIH from controls (Fig. 4k), with AUC=1.0, outperforming plasma cells (AUC=0.89). CD8^+^TOX^+^ T-cells were more abundant on inspection (Fig. 4i). Total CD8+ T-cells were also discriminating (AUC=0.97), and superior to plasma cell enrichment, but with wider variability and inferior to CD8^+^TOX^+^ enrichment indicating there is both increased infiltration of CD8 T-cells and independently increased skew among this population to the TOX^HI^ state in AIH. CD4^+^FOXP3^+^ T_reg_ enrichment favored AIH, but contrary to snRNA-seq prediction was inferior to plasma cell prediction in distinguishing AIH (AUC=0.83) (Figs. 4l,m,n).

AIH tends to have higher injury (and ALT) than MASH for example, raising the possibility that CD8^+^TOX^+^ T-cells simply mark degree of injury. In this study however, ALT spanned a wide range of values across diagnoses, and poorly correlated with CD8^+^TOX^+^ T-cell area density, indicating CD8^+^TOX^+^ T-cell is not a generic marker of hepatocyte injury (Fig. 5o).

**Figure 5.**
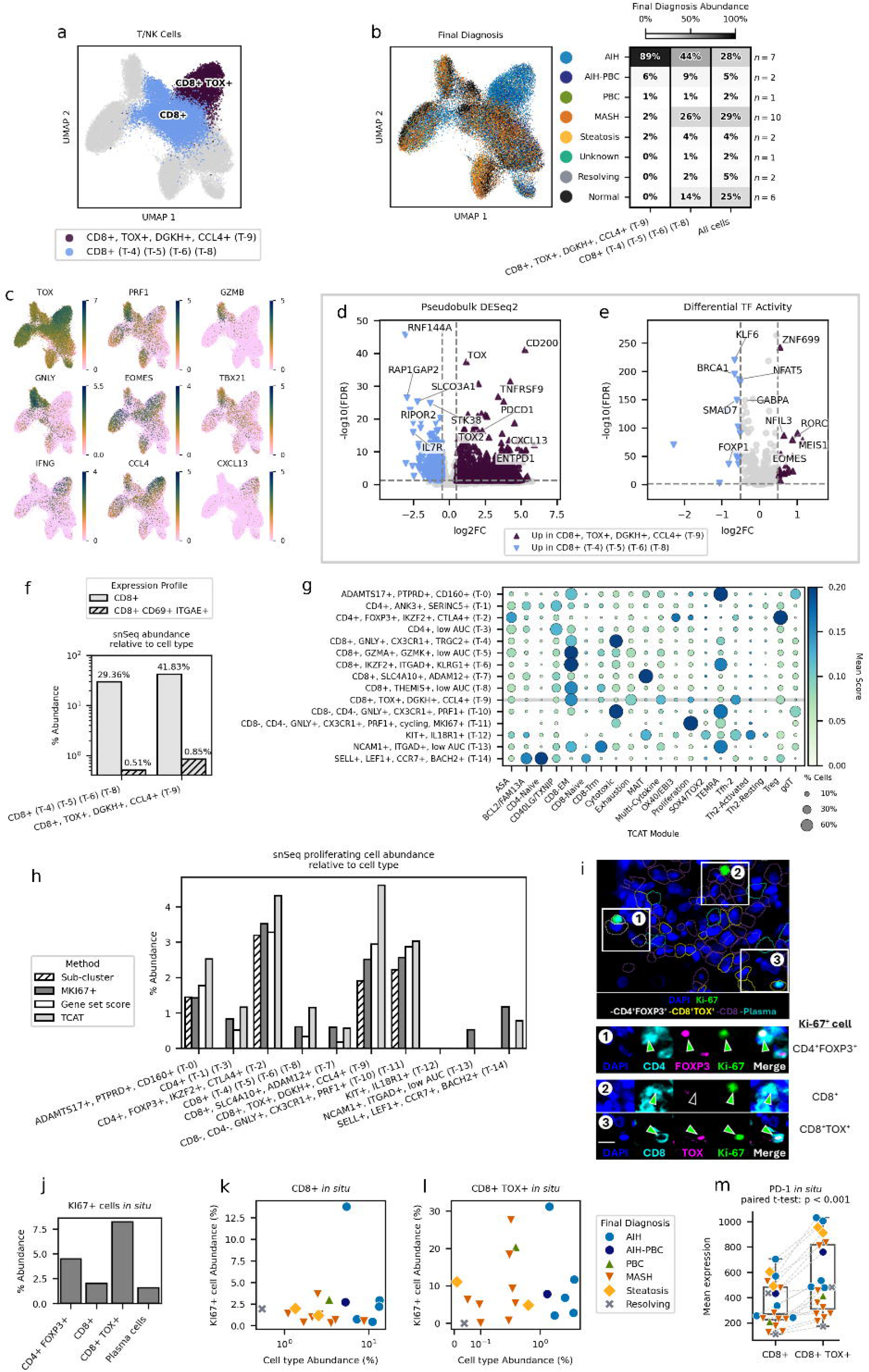
Phenotyping CD8+TOX+ T-cells. (a) UMAP of T/NK-cells denoting T-9 (CD8^+^TOX^+^ T-cells) versus other CD8+ populations (T-4, T-5, T-6, and T-8), excluding MAIT-cells which defines the relevant populations for subsequent analyses, including differential expression. (b) UMAP of T-cells colored by diagnosis; accompanying table shows the fractional abundance of cluster by diagnosis per CD8^+^ T-cell type (CD8^+^TOX^+^ T-cells versus other). (c) Select genes as UMAP heatmaps specifying specific axes of T-cell function. d) Volcano plot demonstrating pseudobulk differential expression of CD8^+^TOX^+^ T-cells versus other CD8^+^ T-cells as specified in (a). (e) Volcano plot demonstrating differential TF activity (AUcell and collecTRI) for the same comparison. (f) CD8^+^ T-cell fractions versus CD8^+^CD69^+^ITGAE^+^ fractions. (g) TCAT dotplot as orthogonal representation of putative T-cell function. The x-axis marks select gene modules versus cluster; horizontal gray line highlights CD8^+^TOX^+^ T-cells (T-9). (h) Quantification of proliferation by four methods in snRNA-Seq dataset, see Methods for a description of each. (i) Ki-67^+^ cell counting in situ: machine-learned cell identities were further gated on Ki-67^+^ nuclear staining. Top: outlines show machine-learned cell identity based on the biomarker definitions in Fig. 4f. Bottom: panels are actual staining patterns highlighting colocalization of double- or triple-markers *in situ* to quantify Ki-67^+^ populations. (j) Quantification of Ki-67^+^ populations in situ. (k-n) Cell type abundance, with method in title, for CD8^+^TOX^+^ T-cells (1 and 3) versus CD8^+^ T-cells (2 and 4) plotted against the abundance of proliferating cells, by individual patient/diagnosis.

Collectively, these results affirm CD8^+^TOX^+^ T-cell enrichment as the most distinguishing cell type of AIH and provide a pathway toward improved diagnosis using conventional immunohistochemical tools.

### Understanding the identity and function of CD8^+^TOX^+^ T-cells

We next sought to better understand the putative identity and function of CD8^+^TOX^+^ T-cells. Using the snRNA-seq data, we focused on cluster T-9, which had representation from most diseased samples, but primarily comprised cells from AIH patients (Fig. 5a,b). TOX is a transcription factor that governs both activation and exhaustion of T-cells, and is typically associated with chronic, antigen-specific activation^61–64^. Among CD8 T-cells, this population expressed intermediate/low cytotoxicity markers compared to other populations like clusters T-4 (*GNLY*, *PRF1*), T-5 (*GZMA*), and T-10/11 (*GNLY*, *GZMA*, *GZMB*, *PRF1*). Markers of progenitor/memory program were relatively depleted (*TCF7* low, *IL7R* very low, *SLAMF6* intermediate) (Fig. 5d), and tissue-residency markers were only partially represented: ITGAE/*CD103* was upregulated, but without *CD69* or *CXCR6* enrichment. As CD69^+^CD103^+^ T-cells were previously reported to be enriched in AIH^65^, we gated on these markers specifically (Fig. 5f), finding only mild enrichment in CD8⁺TOX⁺ T-cells suggesting these are not the same populations. The activation/exhaustion marker *TOX* was accompanied by *TOX2*, *PD1*, and *EOMES*, and transcription factor (TF) activity analysis (Fig. 5e; Supp. Table 2) corroborated target engagement of EOMES^64,66^, while further highlighting marked upregulation of NR4A2 and NFIL3. The TCAT package^67^ was applied to our T-cell clusters to map the transcriptional profiles of our T-cells to other tumor, infectious, and autoimmune contexts (Fig. 5g), and in particular demonstrated a cytokine-high module including (*IFNG*, *CCL4* and *CXCL13*) in CD8^+^TOX^+^ T-cells (Fig. 5c).

Proliferative capacity and PD1 protein expression were directly measured in this population. Proliferation was queried among T-cell fractions multiple ways in the transcriptomic dataset (see Methods) with surprising concordance (Fig. 5h) and demonstrated substantial proliferative capacity of cluster T-9. This observation was corroborated *in situ* by PhenoCycler evidence of Ki-67 co-positivity (Fig. 5i,j). Both methods suggested CD8^+^TOX^+^ and CD4^+^FOXP3^+^ T-cells were highly proliferative. Evidence of proliferation was found in all AIH and overlap patients (Fig. 5k), whereas other CD8 T-cell proliferation was more variable (Fig. 5l). Consistent with the differential expression result (Fig. 5d), CD8^+^TOX^+^ T-cells demonstrated substantially more PD1 protein expression than CD8^+^ T-cells (Fig. 5m). Detailed sub-clustering and pseudotime trajectory analysis were conducted combining T-9 with the other CD8 populations. These analyses largely affirmed the DE results (Fig. 5d,e) above, with minor differences, but highlighted that there is more a continuum (e.g., *IL7R* to *TOX* or *PDCD1*) than a discrete state transition between T-9 and other T-cells (Supp. Fig. 7).

These observations suggest CD8^+^TOX^+^ T-cells represent a PD1-high, proliferative, terminally differentiated (*IL7R* low) effector population.

## Discussion

AIH remains functionally a clinical diagnosis supported by demographics, history of concomitant autoimmune disease, excluding alternative etiologies, response to therapy, and relapse after therapy withdrawal^6,8–10^. Collectively this makes the diagnosis of AIH syndromic, rather than guided by direct correlates of cellular or molecular pathogenesis. Here we present a cellular taxonomy of AIH contrasted against liver diseases that, before diagnostic biopsy, most closely resemble it. We identify a T-cell marked by CD8 and TOX whose enrichment appears to specifically mark the presence of AIH, and is more discriminating than plasma cells, the historical hallmark cell type of AIH^41–44^. More broadly, our approach establishes a pipeline from snRNA-seq of patient specimens to *in situ* histological markers of liver disease.

Our cohort was assembled prospectively at the point of a clinically-indicated diagnostic liver biopsy. This design allowed us to calculate performance characteristics of the International AIH Group Revised Original AIH Score and the Simplified AIH Score which yielded high sensitivity but modest specificity. The performance characteristics reported herein are remarkably close to those reported by Czaja in previous external validation^8^, indicating our cohort captures the diagnostic uncertainty encountered in real-world practice and serves as a meaningful substrate for asking whether molecular phenotyping can improve discrimination of AIH. Of note, BMI did not differ between MASH and AIH patients in our cohort, underscoring two overlapping clinical realities: lean MASH is a major alternative diagnosis in patients presenting with possible AIH, and that obese patients develop AIH and will be missed if BMI is used as a weighting feature in diagnostic algorithms. Thus, while MASH was the most common alternative diagnosis pre-biopsy, incorporation of BMI or other cardiometabolic comorbidities into AIH scoring systems is unlikely to improve diagnostic performance characteristics.

The first major finding of this work is that CTAs, which comprise the numerical composition of the liver’s parenchymal, non-parenchymal and immune populations, are largely sufficient to distinguish AIH from non-AIH. The discrimination is not perfect, but it exceeds what current syndromic definitions of AIH achieve, and it is obtained without exposing the classifier to any morphological description, serum biomarker, patient sex or demographic. That a purely compositional signature performs this well implies that AIH occupies a distinct cellular landscape in the liver, not merely a distinct inflammatory intensity, an assertion further supported by the lack of correlation between ALT and CD8^+^TOX^+^ T-cell abundance. It is worth highlighting that patient-level clustering by diagnosis was not a given; the clustering could have just as easily separated on fibrosis state, biologic sex, age, or other dominant source of data variation, making it all the more striking that AIH versus non-AIH was the major unbiased axis of variation in cell type distribution.

More striking still was finding that the numerical abundance of CD8⁺TOX⁺T-cells was the most important driver of AIH separation and achieved perfect discrimination in our dataset (AUC = [1.00]). This single marker better discriminated AIH than clustering on CTAs, where all the data were used. Crucially, this prediction of relative enrichment of CD8^+^TOX^+^ T-cells in AIH samples mapped cleanly onto the cell area density *in situ,* offering a direct path from a snRNA-seq observation to a tissue-based measurement that could plausibly inform the diagnosis in the setting where it is needed, the diagnostic biopsy itself.

The transcriptional profile of the CD8⁺TOX⁺ T-cells we describe (*PD1*, *LAG3*, *HAVCR2*/TIM3, *TIGIT*, *ENTPD1*, *TNFRSF9* and *CXCL13*) maps closely onto intratumoral CD8⁺PD1^+^ populations in NSCLC^68^ (Fig. 5d) and melanoma^69^, and onto the EOMES^+^CD8⁺TOX⁺ cells described in HCC^70^, where such cells are clonally expanded, antigen-reactive, proliferative and, in functional assays, show attenuated cytotoxicity. This phenotype of attenuated cytotoxicity with a signature cytokine (CCL4, IFNγ, and CXCL13) profile and enhanced proliferation differs from classic exhaustion where all three functions are attenuated^71^. That said, every marker in the signature including PD1 and TOX themselves is induced by T-cell activation as readily as by chronic stimulation, and the functional readouts across disease contexts conflict. The CD8⁺PD1^+^ cells in murine models of AIH^72^ and autoimmune cholangitis^73^ were strongly cytotoxic, and their depletion protective, suggesting a direct pathogenic role. In contrast, islet-antigen-specific CD8^+^TOX^+^PD1^+^ CD8⁺ cells in human T1DM associate with slower rather than faster islet loss^74^, and consistent with this, inhibiting the exhaustion effector LAG3 in NOD mice reverses this restraint and accelerates autoimmune islet destruction^75^. Whether these cells propagate or attenuate injury in AIH will require direct functional interrogation.

To what degree do the CD8⁺TOX⁺ T-cells match previously identified biology in AIH? As noted, this population overlaps with but is likely not the same as previously described CD69^+^CD103^+^ CD8-T-cells in You *et al* ^65^ which were reported to correlate with AIH disease activity (Fig. 5f). Park *et al* provide helpful nuance, having recently described that murine CD69^+^CD103^+^ CD8^+^ T-cells comprise two distinct populations, with TOX engagement being the hallmark of the chronically antigen-stimulated subtype^76^. We also find MAIT cells express both CD69 and CD103 transcripts and others have reported protein-level expression in liver-resident MAIT cells^76^. Finally, we reviewed with interest Xu *et al*, a study reporting an AIH-only single cell atlas which also contained CITE-seq data^77^. Based on *IFNG*, *CCL4*, *TIGIT*, and *CTLA4* expression among CD8 T-cells we suspect the CD8⁺TOX⁺ T-cells here map to their T-cell clusters T_EM_-e, T_EM_-f, or T_EM_-g, or possibly some combination. Their data was not available in sufficient detail for re-analysis but a request for their data was pending. Interestingly, T_EM_-e,f,g in their study all demonstrated liver-specific clonal expansion, compatible with the active proliferation of CD8^+^TOX^+^ T-cells observed here.

Plasma cells are commonly considered a histological hallmark of AIH^41–44^ and after its recognition as a disease entity, the condition was briefly named plasma cell hepatitis^78,79^. Detailed subsequent phenotyping revealed these cells are neither necessary nor sufficient for a diagnosis of AIH^80^, but in one modern pathology study where viral hepatitis could be more decisively excluded^16^, plasma cells in excess of lymphocytes was the histologic feature with highest predictive value over hepatitis C (70-88%). Interestingly, against a variety of comparator diseases here, the area density of plasma cells had an AUC of 0.90, suggesting that more precise and quantitative plasma cell counting as enabled by multiplexed *in situ* staining may add diagnostic power. Nevertheless, our results suggest quantitation of CD8^+^TOX^+^ T-cells, or even quantifying CD8 T-cells, is superior to plasma cells. Further work is required to understand the heterogeneous role of plasma cells in AIH diagnosis and pathogenesis.

T_regs_ have long been hypothesized to play a mechanistic role in loss of tolerance in AIH. It has previously been reported T_regs_ are depleted in the periphery^81^, and enriched in the liver^31^, the latter observation further solidified by our finding of T_reg_ enrichment by both snRNA-seq and by spatial multiplexed immunofluorescence *in situ*. The precise relationship between T_regs_ in the liver and AIH remains unclear; are these cells insufficient in number, or themselves dysfunctional? Interestingly, TCAT highlighted T_regs_ as having the most prominent antigen-specific activation (“ASA”) profile (Fig. 5g), making their clonal expansion and cognate antigen repertoire of particular interest in future work.

This study also highlighted relative depletion of specific cell types as supporting the diagnosis of AIH, with MAIT cells being among the most predictive. MAIT cells are an innate-like, MR1-restricted lineage that normally accounts for a substantial fraction of the intrahepatic T-cell compartment^82^, and prior work finds robust depletion of MAIT cells in the periphery in AIH patients^28,32^. Reports conflict on whether there is intra-hepatic enrichment^28^ or depletion^32^. We suspect these contradictory reports can be reconciled by quantification method; here MAIT cell abundance was depleted when normalized to total T/NK fraction in snRNA-seq, similar to the CD3^+^ normalization of MAIT cells in Böttcher *et al*^32^. In contrast, anti-Vα7.2 immunostaining of liver sections allows absolute quantification of MAIT cells per unit area and was reported to be elevated in AIH^28^. Thus, there is likely an absolute increase in intra-hepatic MAIT cells, but a relative decrease driven by profound expansion of other T-cell populations.

This study has important limitations. First, because patients were recruited prospectively, before their diagnoses were known, the number of ultimately confirmed AIH cases is modest (n=6 pure, n=8 including overlap). This number is itself informative: it reflects the difficulty of predicting AIH pre-biopsy and argues that future studies will need to enrich for AIH biology through post-biopsy recruitment or multi-center collaboration. The converse benefit is that our control population is diverse and captures many of the actual alternative diagnoses that clinicians must discriminate from AIH, which strengthens rather than weakens any claim of AIH-specific biology. Another cohort specific caveat is clinical under-utilization of anti-soluble liver antigen (SLA) at our center. Anti-SLA is a highly specific auto-antibody^83^ in about 21%^84^ of AIH patients, but was only drawn in 6 of 26 individuals in the diagnostic cohort. That said, anti-SLA was negative in the 3 AIH cases it was checked, and the primary drawback of current diagnostic strategies were false positives, so more complete anti-SLA data would not have resolved the incomplete diagnostic picture highlighted in this prospective study.

The most notable individual gap in our cohort is the under-representation of drug-induced liver injury (DILI). DILI is widely regarded as the most difficult of AIH’s mimics to exclude clinically and histologically, yet it was infrequent in our prospectively enrolled cohort. This reflects, we suspect, a clinical pattern in which patients with a strong suspicion of DILI are more often observed off the suspected offending agent than biopsied. DILI is also itself a heterogeneous disorder likely containing autoimmune-like conditions^39,40,85,86^. Further work is therefore needed to determine whether CD8⁺TOX⁺ T-cells mark AIH globally or whether some classes of DILI will require an alternative approach to discriminate.

A final limitation concerns the scope of our *in situ* validation which was performed on the same patients from whom snRNA-seq was obtained. These experiments were designed to determine whether snRNA-seq cell-type abundances are recoverable by *in situ* measurement in a way that is quantitatively meaningful and diagnostically relevant. This was not a given, and our data answer this question clearly, and in the affirmative, establishing a pipeline that can now be applied to the independent validation cohort that must follow. We have further specified a ranked “differential diagnosis” of other cell types that might distinguish AIH alone or in combination with other markers (Fig. 3), with substantial further work required to optimize *in situ* detection of more obscure cell populations like mRegDCs, or proliferating cDC1 populations.

Taken together, our findings nominate CD8⁺TOX⁺ T-cell abundance as a candidate tissue biomarker for AIH, taking a concrete step toward a rational, positive diagnosis of a disease currently identified by exclusion. These data also provide a critical cellular clue for understanding AIH mechanism. Although the disease responds to broad immunosuppression, implicating a defect in tolerance, no consensus autoantigen has been established, no effector cell type has been mechanistically defined, and the choice among emerging targeted immunomodulators remains empirical. A specific, enriched, putatively antigen-experienced T-cell population offers a tractable starting point for each of these gaps. The pipeline established here, from unbiased snRNA-seq of diagnostically ambiguous biopsies to *in situ* validation of candidate cell types, is generalizable and offers a path toward improved diagnosis of AIH.

## Acknowledgments

The authors would like to thank our patients for their participation in this work, without which none of this would have been possible. Similarly, to the interventional radiology faculty at Massachusetts General Hospital, and in particular Julie Mucciarone, Melissa Chittle, and Meredith Preziosi, whose collaboration with our research team enabled timely collection of most of the research samples in this study. We thank the entire MGH Hepatology faculty, and many fellows who contributed referrals for this project. The authors would like to acknowledge Dr. John Vierling for his scientific inpfut and enthusiasm for this work, and Suzan Dijkstra for her careful reading of the manuscript and her input on the candidate function of CD8⁺TOX⁺ T-cells. The authors are grateful for Alan Mullen for his advice, manuscript feedback, and for allowing the authors to piggyback on an IRB-approved protocol he helped maintain.

## Methods

### Study Design

Patients were enrolled under an Institutional Review Board (IRB)-approved protocol (Massachusetts General Hospital) allowing the collection of liver tissue and peripheral blood at the time of clinically indicated liver biopsy. Liver samples were obtained either from percutaneous diagnostic biopsies or from excess surgical tissue following resection procedures. All patient samples were anonymized at the time of collection and assigned a unique study identifier. A coded key linking identifiers to clinical information was maintained in a password-protected spreadsheet stored on an encrypted institutional hard drive.

Patients were prospectively recruited at Massachusetts General Hospital, a major tertiary care referral center in the Northeast for autoimmune and cholestatic liver diseases. Potential participants were identified at the time of referral for clinical liver biopsy, if the treating hepatologist considered AIH to be among the top two diagnostic considerations. Patients were required to have negative serological testing for hepatitis A, B, and C. Patients with greater than 3 standard alcohol beverages per day, or on therapy with corticosteroids (≥5mg prednisone or equivalent), azathioprine, or mycophenolate in the prior 6 months were excluded. Established diagnoses of PBC were not explicitly excluded if the intention of the biopsy was to rule out AIH or AIH overlap.

Normal liver tissue was obtained from patients undergoing hepatic surgical resection for benign or malignant conditions. These individuals were required to have no history of underlying liver disease aside from hepatic metastases. Additional exclusion criteria for this control group included aminotransferase levels >40 IU/L and recent systemic chemotherapy within six months of tissue collection.

For percutaneous biopsies, tissue was collected directly from the biopsy needle into pre-chilled Belzer UW transplant preservation media (Bridge to Life, Madison, WI) and washed briefly. Biopsies were then gently dried on sterile plastic and divided into thirds, with each portion placed into pre-labeled cryovials and slow-frozen by burying in dry ice. From biopsy acquisition to dry ice was < 2 minutes. In surgical cases, liver resection material was first processed by a clinical pathologist to identify resection margins; only grossly uninvolved healthy liver tissue margins were used. The tissue was subsequently washed in PBS, transferred to UW transplant preservation on ice, and then divided into rice size pieces also adhered to cryotubes and slow-frozen as above. From patient resection to dry ice was <1h. Frozen tissue was maintained in liquid nitrogen storage until tissue processing.

### Tissue processing and single nuclei suspension

Both percutaneous and surgical frozen liver samples were processed for snRNA-seq using a protocol adapted from Drokhylyansky et al^87^. Frozen cryovials containing one third of a liver biopsy (approximately 0.65cm x 1mm cylinders; <2mg wet weight) were partially thawed in gloved hands, and transferred to cold Tween, Salts, Tris buffer (TST^87^ plus 0.5% bovine serum albumin (BSA)) on ice. Tissue fragments were finely minced using sterile scissors in a chilled Eppendorf tube, then transferred via wide-bore pipette to a dounce homogenizer. Homogenization was performed using 20 strokes each with pestle A and pestle B. A 10 µL aliquot of the homogenate was mixed with trypan blue and assessed under by light microscopy to evaluate nuclear yield, quality and cell lysis. In rare instances where intact cells were still present, additional chemical lysis was allowed for 3-10 minutes, before resampling and reimaging. Most samples required no delay and proceeded immediately.

The homogenate was filtered through a 40 µm cell strainer into a 15 mL conical tube containing Salts and Tris (ST) buffer^87^ with 1% BSA, and centrifuged at 500 g for 5 minutes at 4°C. The nuclear pellet was resuspended in 40 µL of recovery buffer and filtered again through a 10 µm mini-strainer (Pluriselect). Final nuclei counts were obtained using a manual hemocytometer, with a goal to load 10,000 nuclei per sample for downstream processing. Single-nuclei suspensions were then loaded on a Chromium controller (10X Genomics). To preserve sample quality, samples were processed in <1 hour between thawing and chip loading, which limited processing to a maximum of two samples at a time. All samples were prepared from a single lot of 10x kits and processed over a one-month period in March of 2023.

### snRNA-seq library preparation and sequencing

Nuclei were processed according to the manufacturer’s protocol for the 10x Genomics Chromium Single Cell 3’ v3.1 platform with the following choices or modifications. cDNA quality control was conducted with an Agilent TapeStation, and samples with poor quality at this step were not carried forward. Where available, samples were re-processed. In the cDNA amplification step, two additional cycles were used based on a 10X application note for snRNA-seq. Libraries were sequenced on Illumina NovaSeq platforms using GeneWiz. Libraries were sequenced to a target depth of 500 million reads per sample (corresponding to ∼50,000 reads per nucleus). Sequencing data were demultiplexed and processed through the 10x Genomics Cloud Analysis Portal, including FASTQ generation and Cell Ranger count matrix generation with default settings.

### snRNA-seq pre-processing

#### snRNA-seq quality control

CellBender v0.2.084^88^ was run on each sample individually for ambient RNA removal (Supp. Fig 2a). Most samples were run with parameters estimated by CellBender, however some samples failed with estimated parameters. Samples that failed were run on specified parameters based on the estimated values but changed to meet the CellBender requirements (Supp. Table 3). Post-ambient-removal cells were stored in the ‘cellbender’ layer of the anndata object and pre-ambient removal cells were stored in the .X layer. Raw counts for pre and post-ambient removal are stored in counts and cb_counts layers, respectively.

Scanpy v1.10.285^89^ on Python v3.9.0 was used for data processing and analysis. Quality control steps are shown in Supp. Fig. 2b. In the .X layer: genes that existed in fewer than three cells, and cells that contained fewer than 200 genes in a sample were removed. In the cellbender layer: cells where the percentage of mitochondrial genes exceeded 20% of total genes were removed. Highly variable genes were calculated for each sample using sc.pp.highly_variable_genes() function with the following parameters: min_mean=0.0125, max_mean=3, min_disp=0.5, batch_key=‘sample’, layer=‘cellbender’. Highly variable genes were selected if at least n samples shared the highly variable gene, where n>2. n was selected by which n brought the number of highly variable genes closest to 4000.

Both the .X and the ‘cellbender’ layer were normalized with sc.pp.normalize_total function with parameters: target_sum=10000, exclude_highly_expressed=True, key_added=‘norm’, layer_norm=‘after’ and then sc.pp.log1p() function with default parameters. Then, Scrublet v0.2.386^90^ was run for doublet detection on the .X layer data using default parameters. PCA was performed with the scanpy.tl.pca() function on the cellbender layer using default parameters. Batch-balanced k-nearest neighbors (BBKNN)^91^ was run for batch correction with scanpy.external.pp.bbknn() function using default parameters (Supp. Fig 2c).

#### Annotation

To identify cell clusters, the sc.tl.leiden() function was run with varying resolution. Resolution for each group of cells was determined by biological relevance and whether the resulting clusters were well defined by differentially expressed genes. Differential expression was performed with scanpy.tl.rank_genes_groups() function and parameters: method=’overestimated_var_t_test’. Marker genes were determined by a combination of cluster specificity and literature-based relevance for the gene and cell type of interest.

Doublets were identified and removed based on a combination of a high doublet probability score from scrublet and/or the clustered separately from definite singlets and contained marker genes characteristic of two or more cell types (Supp. Fig. 8).

To cluster subtypes, cells were grouped by coarse cluster labels (Figure 1c). Highly variable genes, PCA, batch correction, UMAP and leiden were recalculated following the same functions and steps used for the all-cell group. T/NK Cells (C-5) and cholangiocytes (C-2) sub-clustering used the .X layer. Myeloid cells (C-4), endothelial cells (C-3), B-cells (C-6), hepatocytes (C-0) and hepatic stellate cells (C-1) used the ‘cellbender’ layer. Layer usage depended on subjective judgement of whether the clustering of a sub-type produced technical effect or biological noise.

### Definition and analysis of cell type abundances (CTAs)

A cell-type abundance (CTA) vector (*Γ^S^*) is the compositional representation of a sample *S*, whose elements correspond to the relative abundances of cell types *j* in parent population *P* and sum to one. CTA vectors can be defined at any level of the cellular hierarchy by normalizing counts within the relevant parent (*P*) population.

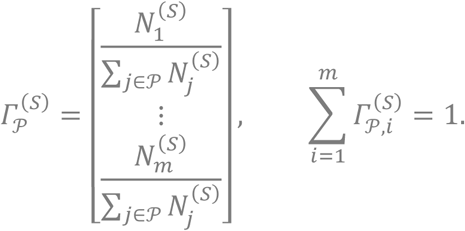

When *P* is the coarse cell types, then for each sample the CTA vector is composed element-wise of the fractional abundance of hepatocytes, T/NK cells, cholangiocyte, and so on, where each fractional abundance of cell type *j* is the total number of cell type *N_j_* divided by the total cells in that parent population (denominator summand). In the case of a cellular subtype, like myeloid cells, the same logic holds but the summand would equal the number of myeloid cells in that sample, and N_j_ corresponds to the numbers of cellular subtypes among myeloid cells.

Composite CTAs were produced by concatenating (stacking) levels; for example, the composite coarse plus immune CTA for Fig. 3 comprised coarse, lymphoid, myeloid and B/plasma CTAs:

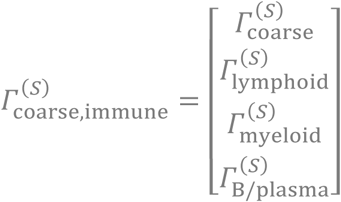

For sample (patient) level clustering, a sample by CTA matrix **Γ** was defined, where *S* denotes all samples in the snRNA-seq dataset:

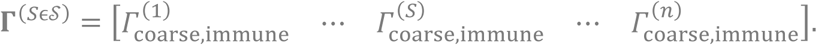

Initially, **Γ**(SɛS) was input for clustering. However, this proved to be problematic as more abundant species, like hepatocytes which represented relative abundances as high as 80 to 90% in some samples, dominated PCA space and variance. Therefore, each row entry of **Γ**(SɛS) (fractional abundance of a given cell type, or sub-cell type) was normalized by the sum across that row, for all rows (Γ̃). Thus, lowly abundant species that varied significantly by sample (patient) could contribute to variance in ordinal space, whereas relative abundances that varied little between samples would contribute little to PCA space.

### CTA Clustering and PCA

Unsupervised clustering was performed on this normalized, combined coarse and immune CTA matrix Γ̃ (Fig. 3c) using sklearn.cluster.SpectralClustering() with default parameters, sklearn.cluster.AgglomerativeClustering() with default parameters, and sklearn.cluster.KMeans() with default parameters. K-means was run for 5000 iterations on different random seeds with all samples. The n_clusters parameter was determined by combination of a consistency metric, here we used the Jaccard index, and biological interpretation in the results (Supp. Fig. 5).

To inform the selection of the number of clusters, the average pairwise Jaccard Index was calculated for each clustering method with and without bootstrapping. Given a clustering algorithm, and a cluster size c: A reference clustering was generated with all samples and c clusters. Then, for n iterations data was subsampled by sample_id (n=25 for bootstrapping, n=31 for all samples), clustered into c clusters (with a varying random seed), and the Jaccard index was calculated with respect to the reference clustering. The mean was taken between all scores for a given c. Shared cluster identity was determined by whether samples S4, S22, or S47 were shared.

For each clustering algorithm the average Jaccard Index was calculated for cluster sizes between 2 and 8. For spectral, agglomerative, and k-means the average was taken across 100, 100 and 500 iterations, respectively.

### CTA Regression

A logistic regression model was trained using Γ̃ as input and binary disease diagnoses as labels (determined by final diagnosis in the metadata) using scikit-learn v1.5.1 sklearn.linear_model.LogisticRegression() function with parameters: random_state=0, multiclass=’auto’. The binary labels (Fig. 3g) used for comparison were: AIH Exclusive, AIH Inclusive, AIH Overlap Excluded, AIH vs MASH, AIH vs Disease.

### Differential expression analysis

Samples were pseudobulked with pegasuspy^92^ v1.10.2 pegasus.pseudobulk() function with parameters: groupby=’sample_id’, mat_key=’cb_counts’ (here, ‘cb_counts’ refers to the CellBender raw counts). Samples were only kept in a pseudobulked cell type if the sample had at least 10 cells in the cell type, and a cell type was included if it was represented by at least 3 samples.

Differential expression was run with pydeseq2 v0.4.12. Pseudobulked cell types were stacked into one dataframe to create the pydeseq2.dds.DeseqDataSet() object with parameters: design_factors=[‘celltype’], refit_cooks=True^93^. The function deseq2() was run on the DeseqDataSet object with default parameters (Supp. Table. 1).

### Cell types with highest expression of gene

For every gene measured, the coarse cell type with the highest mean expression compared to other coarse cell types was identified for the following groups: All cells, and each final diagnosis group (Supp. Table. 4).

### Transcription Factor Activity Prediction

Transcription factor activities were inferred with decoupler v1.8.0 ^94^ decoupler.run_aucell()^95^ with parameters: seed=42,min_n=5,seed=42,use_raw=False. The collecTRI regulon network^96^ was referenced with function decoupler.get_collectri() with parameters: organism=’human’. The input expression data was the T/NK cells (C-5) ‘cellbender’ layer only including genes that were most highly expressed in T/NK cells (C-5) in any diagnostic group (see above).

To calculate differential activity between (T-9) and (T-4, T-5, T-6, T-8), a two-sided Wilcoxon rank sum test was performed between cell-resolution AUCell scores with function scipy.stats.mannwhitneyu() and parameters: alternative=”two-sided”. P-values were adjusted with statsmodels.stats.multitest.multipletests() and parameters: method=”fdr_bh” from statsmodels v0.14.6 (Supp. Table 2).

### StarCAT

StarCAT v1.0.10^67^ function starCAT.fit_transform() with default parameters was run on T/NK cells (C-5) raw CellBender counts layer=‘cb_counts’ with the TCAT.V1 reference.

### snRNA-seq Proliferating Cell Identification

Subclustering, TCAT (Methods: StarCAT), and two score-based methods were used to identify proliferating cells in snRNA-seq data.

The subclustering approach was to subcluster down to lineages, or to individual subcell types to investigate whether a proliferating cluster separated from the main body of each cell type marked by canonical cycling genes (Supp. Fig. 9a). Harmony v0.2.0^97^ (with Python v3.12.0 and Scanpy v1.12) was used for batch correction since it was able to handle groups of cells where some samples had low representation. Major T/NK subtypes [(T-9), (T-4, T-5, T-6, T-8), (T-7), (T-1, T-3), (T-2), (T-10, T-11), (T-0), (T-12), (T-13), (T-14)] were isolated for sub-clustering: sc.pp.highly_variable_genes() with parameters: min_mean=0.0125, max_mean=3, min_disp=0.5, batch_key=’sample_id’,layer=’cellbender’; scanpy.tl.pca() with parameters: layer=’cellbender’; harmonypy.run_harmony() with parameters: adata.obsm[’X_pca’], adata.obs, ’sample_id’, max_iter_harmony=50; scanpy.pp.neighbors() with parameters: n_pcs=30, use_rep=“X_pca_harmony”; scanpy.tl.umap() with default parameters, and scanpy.tl.leiden() with a varying resolution parameter.

TCAT-inferred proliferating cells were identified with the module “Proliferation Binary” (Supp. Fig. 9b).

Scoring methods included scanpy.tl.score_genes_cell_cycle() with S and G2M gene sets from Tirosh et al.^98^; and MKI67 expression (Supp. Fig. 9c,d,e). Cutoffs for each scoring method were subjectively determined based on visual inspection of scoring distribution (S=0.2, G2M=0.2, MKI67= 0) (Supp. Fig. 9f-j).

### Spatial proteomic tissue processing and imaging

Cell types distinguishing AIH from non-AIH (Fig. 3h) were prioritized for *in situ* area density measurement using spatial proteomics (Akoya PhenoCycler), and was designed to capture CD8^+^ T-cells, CD8^+^TOX^+^ T-cells, CD4^+^FOXP3^+^ T-cells, plasma cells, cDC1, cDC2 and proliferation markers. A panel of viable primary antibodies were screened on two AIH and two normal tissue formalin-fixed paraffin embedded (FFPE) slides sequentially using Multiple Iterative Labeling by Antibody Neodeposition using an established protocol^99^, notably using conventional primary and secondary antibodies. A second round of prototyping was conducted on the PhenoCycler and revealed poor signal-to-noise for both markers required for identification of cDC2 (Table 4). An optimal combination was identified for the remainder of cell types and conjugated antibodies for each primary were ordered from Akoya, custom conjugating as necessary. The final 15-plex panel (Table 4) was processed and imaged on a PhenoCycler Fusion.

**Table 4.**
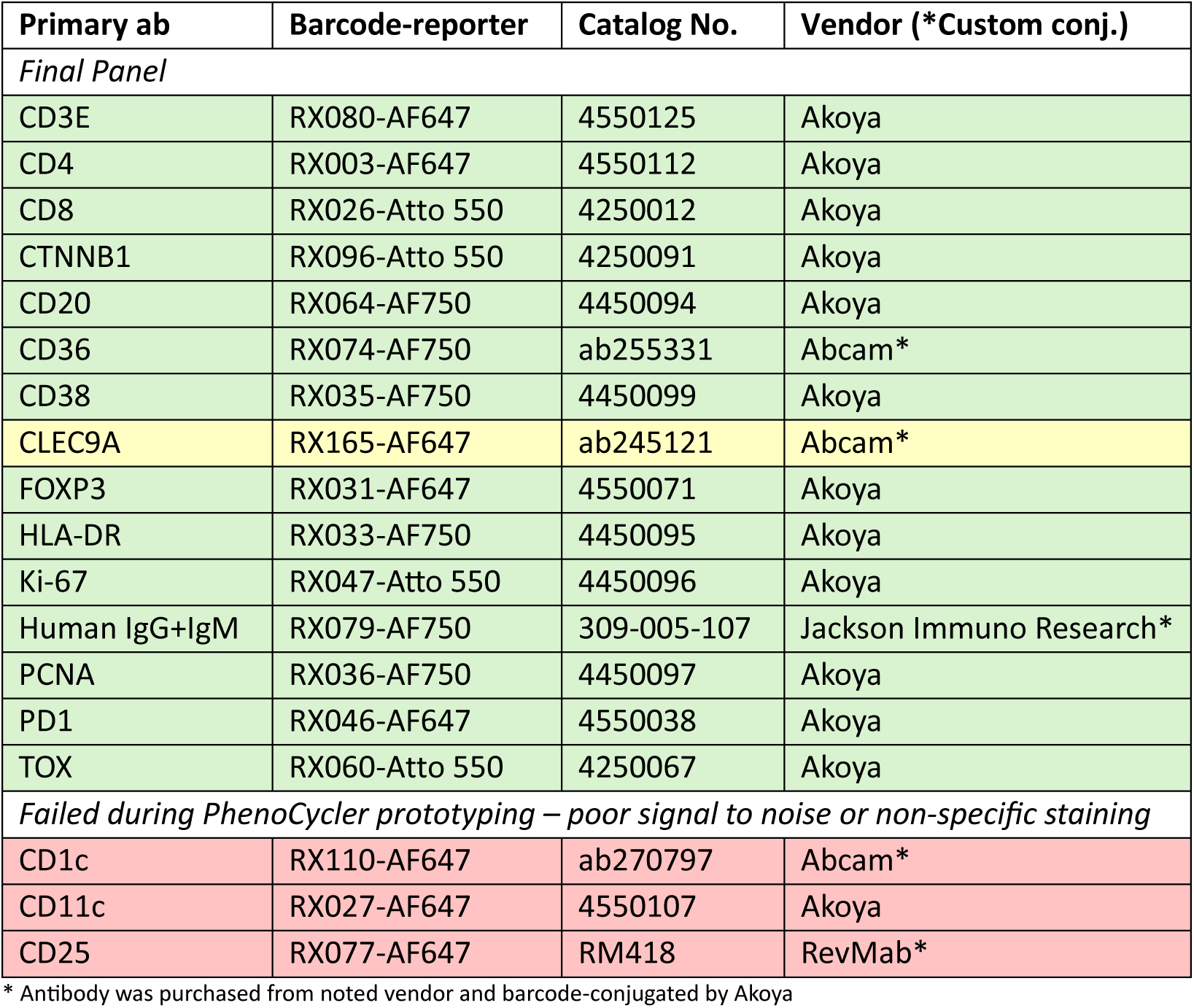

| Primary ab | Barcode-reporter | Catalog No. | Vendor (*Custom conj.) |
| --- | --- | --- | --- |
| *Final Panel* |  |  |  |
| CD3E | RX080-AF647 | 4550125 | Akoya |
| CD4 | RX003-AF647 | 4550112 | Akoya |
| CD8 | RX026-Atto 550 | 4250012 | Akoya |
| CTNNB1 | RX096-Atto 550 | 4250091 | Akoya |
| CD20 | RX064-AF750 | 4450094 | Akoya |
| CD36 | RX074-AF750 | ab255331 | Abcam* |
| CD38 | RX035-AF750 | 4450099 | Akoya |
| CLEC9A | RX165-AF647 | ab245121 | Abcam* |
| FOXP3 | RX031-AF647 | 4550071 | Akoya |
| HLA-DR | RX033-AF750 | 4450095 | Akoya |
| Ki-67 | RX047-Atto 550 | 4450096 | Akoya |
| Human IgG+IgM | RX079-AF750 | 309-005-107 | Jackson Immuno Research* |
| PCNA | RX036-AF750 | 4450097 | Akoya |
| PD1 | RX046-AF647 | 4550038 | Akoya |
| TOX | RX060-Atto 550 | 4250067 | Akoya |
| *Failed during PhenoCycler prototyping – poor signal to noise or non-specific staining* |  |  |  |
| CD1c | RX110-AF647 | ab270797 | Abcam* |
| CD11c | RX027-AF647 | 4550107 | Akoya |
| CD25 | RX077-AF647 | RM418 | RevMab* |

\* Antibody was purchased from noted vendor and barcode-conjugated by Akoya

Slides from a subset of the diagnostic cohort (all AIH and overlap, MASH/steatosis, resolving and PBC, n=23) were obtained, with 2 patient sections per slide, not organized by diagnosis to minimize batch effects. Slides were photobleached for 8 hours, then PhenoCycler Fusion was run across 6 rounds for each slide by the Dana Farber Cancer Institute (DFCI) Pathology Core. Staining quality was reviewed by author MS and the DFCI Pathology senior staff analyst after each run; slides were re-run as necessary. Subsequently, careful review of staining quality for each slide and each marker, blinded to diagnosis, revealed subjectively successful staining with a few minor exceptions. Three samples were excluded due to artifact; one sample was rerun twice, without resolution, containing two samples both of which suffered the same artifact (very high autofluorescence, nonspecific staining of most markers); this slide was excluded. The third was a singleton and appeared to be lifting off the slide; using a new slide did not resolve this issue. The final study (n=20) contained AIH (n=5), AIH-PBC overlap (n=1), MASH (n=9), steatosis (n=2), resolving (n=2), and PBC (n=1). In addition, staining for CLEC9A, which worked in MILAN prototyping, showed off-target hepatocyte nuclear staining in a little more than half the samples using the barcoded/conjugated Akoya primary, thus preventing localization of cDC1 across the whole cohort. The main analysis therefore focused on cell types for which at least two markers were present to validate cell type identity (CD8^+^ T-cells, CD8^+^TOX^+^ T-cells, CD4^+^FOXP3^+^ T-cells, and plasma cells).

### Spatial proteomic data analysis

#### Segmentation and single cell measurement

Cells were segmented using Instanseg^60^ in QuPath^100^ using the layers expected to cleanly define cell shape, and omitting markers that might have non-specific patterns (IgG+IgM), poor signal to noise (PD1), or represented subsets of cell function (Ki67, PCNA, FOXP3, TOX): DAPI, CD3, CD4, CD8, CD20, CD38, and CTNNB1. Fluorescence measurements per-cell were calculated in Qupath including mean, median, min, max, and standard deviation.

#### Annotation of single cells in spatial proteomic data

Cell types were defined using the marker panel logic in Fig. 4f, and subjectively confirmed to be of the expected morphology for each cell type. Blinded to sample diagnosis and ID, canonical examples of double positive cells were annotated by hand across both samples of each slide, and then a machine learning classifier was applied to annotations to learn the rest of the data. Possibly due to batch effect, a model trained on one slide transferred imperfectly to the next slide. Therefore, the first few slides (in random order) were combined for a master model, and then subsequently the master model was applied to each new slide, and then fine-tuned iteratively on each subsequent slide by correcting algorithmic mistakes. When roughly <1 in 15 cells required annotation correction, the fine-tuning was considered sufficient.

#### Quantification of cell type-specific area density

The entire liver biopsy region was outlined, and very large empty areas were excluded (vessels with diameter > 100µm), for each sample. From these, the total area per sample was calculated in units of millimeter (mm). Using whole-specimen annotations of machine-learned CD8^+^ T-cells, CD8^+^TOX^+^ T-cells, CD4^+^FOXP3^+^ T-cells, and plasma cells, each cell type count was divided by the tissue area to obtain the per-patient area density for each cell type.

#### Quantification of Ki-67 positivity from spatial proteomic data

The QuPath data matrix was exported as a tab delimited table comprising cells (rows) by fluorescence measurements, and also contained row-wise metadata (sample, diagnosis, slide, annotation, etc). These data were preprocessed in python to remove cells with measurements more than 75% composed of NaNs, and then mapped to anndata^101^ structure where the obs were cells, the vars were quantitative measurements (median, mean, min, max, and standard deviation) of cell structures (Cell, Nucleus, Membrane, Cytoplasm) as well as cell shape measurements. Metadata was saved in .obsm and .varm. Cells were filtered to only include classified cells (CD8+, CD8+ TOX+, CD4+ FOXP3+, Plasma Cells), and quantitative measurements were filtered to exclude any “Nucleus” measurements as well as any “Min” or “Mean” measurements. In Scanpy v1.12, data was log-transformed using scanpy.pp.log1p() with default parameters and batch corrected using Harmony v0.2.0 harmonypy.run_harmony() with parameters: data=adata.X, batch=”sample_id”, max_iter=50 (Supp. Fig. 10a,b). Batch-corrected data was then clustered with: scanpy.pp.pca() with default parameters; scanpy.pp.neighbors() with parameters: n_pcs=30; scanpy.tl.umap() with default parameters; and scanpy.tl.leiden with parameters: resolution=0.2, n_iterations=3 (Supp. Fig. 10c). Ki-67 high clusters were observed (Supp. Fig. 10d) and specified after sub-clustering (Supp. Fig. 10e-i).

## Author contributions

Conceptualization: MSS

Methodology: MSS, DMS, MFT

Investigation: MSS, DMS, SWK

Visualization: MSS, DMS

Funding acquisition: MS, WG

Project administration: MS, GB, GML, WG

Supervision: GML, ACV, WG

Writing – original draft: MSS, DMS, WG

Writing – review & editing: MS, DMS, WG

