## Supplementary Information for "Hepatic CD8^+^TOX^+^ T-cells are a hallmark of autoimmune hepatitis"

### Annotation of snRNA-seq cell types

*Hepatocytes*

As expected, hepatocytes were abundant in snRNA-seq samples and marked by expression of the lineage-defining transcription factor *HNF4A*, complement (*C5*, *C6*, *C8A*), fibrinogen (*FGG*, *FGA*), and coagulation factors (*F5*, *F9*, *F10*) among others. The hepatocyte cluster demonstrated a zonal gradient (Zone 1: *PCK1*, *GLS2*; Zone 3: *CYP2E1* and *GLUL*) (Supp. Fig. 3a).

*Lymphoid lineage (continued)*

Cells in this population were split into CD8-high (T-[4,5,6,7,8,9,13]), CD4-high (T-[1,2,3,14]), and CD3-low, CD4-low (T-[0,10,11,12]). CD8 populations variously represented a cytotoxic fraction (*GNLY*-high, T-4; *GZMA*-high, T-5), an *IKZF2*/Helios fraction (T-6), and an activated/exhausted fraction (T-9; *LAG3*, *TOX*, *PD1*)[3,4]. In the CD4 fraction, naive/memory T-cells (T-14) were differentiated by expression of lymphoid homing and Wnt signaling genes (*CCR7*, *SELL*, *LEF1*). Putative T-regulatory ($T_{reg}$) cells (T-2) were identified by their enrichment of *CD4*, *FOXP3*, *IKZF2*, *IL2RA* (CD25), and *CTLA4*[5,6]. Among CD4 and CD8 depleted populations, T-12 was marked by *KIT* and *IL18R1* suggestive of an innate lymphoid cell (ILC). The clusters T-0 and T-10 likely capture a mixture γδ-T-cell (*TRDC*) and NK markers (*NCR1*, *NCAM1*) as well as expression of killer lectin-like receptors (*KLRF1*, *KLRB1*, *KLRD1*, *KLRC*). T-0 was distinguished by non-canonical markers (*ADAMTS17*, *PTPRD*) while T-10 was particularly high in cytotoxic markers (*PRF1*, *GNLY*, *GZMB*), and the homing receptor *CX3CR1*, these latter categories shared with the CD8 expressing population T-4.

*B-/plasma cells*

Plasma cell enrichment is strongly associated with AIH[7–10], while B-cell depletion is therapeutic in refractory AIH[11,12]. Both cell types were well represented in snRNA-seq as mature naïve B-cells (B-0, B-1) and plasma cells (B-2) (Fig. 2g). Mature naïve B-cells were distinguished by expression of IgD (*IGHD*), IgM (*IGHM*), antigen processing and presentation (*HLA-DRB1*, *HLA-DRA*, *WDFY4*), and B-cell antigen receptor CD20 (*MS4A1*)[13,14] (Fig. 2h,i). Plasma cells were defined by expression of a variety of immunoglobulin heavy chain classes (*IGHG1*, *IGHG3*, *IGHG4*, *IGHA1*), lambda and kappa chain constant regions (*IGKC*, *IGLC2*), adhesion molecule *CD38*, and plasma cell differentiation regulator *PRDM1*[14,15].

*Myeloid lineage (continued)*

Kupffer cells are stationary liver resident macrophages that are found within the hepatic sinusoids, and were identified by enrichment of *MARCO*, *VSIG4*, and *CD163*, as well as depletion of *SIRPA*[16] (Fig. 2e,f). Within the Kupffer cell population were a group of cycling cells (M-2) marked by *MKI67* and *DIAPH3*. Monocyte-derived macrophages migrate to the site of injury and can be profibrogenic as well as involved in fibrosis recovery; they are distinguished from Kupffer cells by expression of *CPM*, *DHRS9*, and *GPNMB*[17,18]. Within the monocyte-derived macrophages was a population with both Kupffer and monocyte-derived markers (M-3), a phenotype associated with Kupffer cell depletion in MASH[19]. Putative monocytes were identified by enrichment of *VCAN*, *FCN1*, and *LYST*; they were clustered into quiescent (M-6) and activated (M-13) populations. Plasmacytoid dendritic cells (pDCs), which initially clustered with B-cell/plasma cells as expected, were co-clustered with myeloid cells where they remained discrete but closest on the UMAP to cDC2, and were marked by *TCF4*, *CUX2*, and *ZFAT*[20,21] (Fig. 2e,f).

*Endothelial Cells*

The liver receives oxygenated blood through the hepatic artery and nutrient-rich blood from the intestines through the portal vein; this blood then flows through fenestrated sinusoids lined with liver sinusoidal endothelial cells (LSECs) into the central vein. LSECs (E-2) can be differentiated from other liver endothelial cells (LECs) by expression of scavenger and mannose receptors (*STAB1*, *STAB2*, *MRC1*) as well as a lack of endothelial cell adhesion molecule *PECAM1*[22–24]. LECs lining the portal vein and hepatic artery (E-4) were marked by enrichment of *FLRT2*, *AQP1*, *CD9*, *MECOM*, and *TANC2* [5]. LECs lining the central vein (E-0) were distinguished by increased expression of WNT-signaling proteins (*RSPO3*, *WNT2*, *WNT9B*) as well as *ENG*, *IL1R1*, and *TSHZ2*[23,25,26]. Finally, a population of pericytes (E-5) that line LECs expresses smooth muscle markers (*CHRM3*, *SORBS2*), vascular growth factors (*PDGFB*, *VEGFC*), and *EBF1*[27–29]. A group of cycling LSECs (E-3) was identified by expression of *DIAPH3* and *TOP2A*. A subpopulation of central vein LECs (E-1) exhibit high expression of *TSHZ2*, and enrichment of *PELI2*, *PKHD1L1* and *ARHGAP26* distinctly identifies them.

*Hepatic Fibroblasts and Stellate Cells*

Hepatic fibroblasts and stellate cells are the primary producers of extracellular matrix during liver injury. Here we identify fibroblasts (HS-2), quiescent hepatic stellate cells (HSC) (HS-0), myofibroblasts (HS-1), and a distinct cluster of cells exhibiting a neuronal phenotype (HS-3). HSCs separated into quiescent and activated groups; quiescent HSCs were identified through expression of zonal Wnt signaling protein *RSPO3*, retinol metabolism protein (*RBP1*, *LRAT*), MHC class II complexes (*HLA-DRB1*, *HLA-DRA*), *DCN*, *CYGB*, *GPCR*, and *RELN*[30–32]. Myofibroblasts are differentiated forms of hepatic stellate cells or fibroblasts that are involved in injury repair. Myofibroblasts were identified by their expression of ECM degradation inhibitor *TIMP1*, collagen (*COL1A1*, *COL1A2*), *DCN*, *C7*, and *FBLN5*[32–36]. Portal fibroblasts were defined by expression of components of elastin fibers (*ELN*), *SLIT3*, *NOTCH3*, and *ACTA2*[37,38]. Finally, we identified a cluster exhibiting a neuronal phenotype (HS-3) defined by *NRXN1*, *GRIK2*, *NRXN3*, and *ZNF536*.

*Cholangiocytes*

Inter-hepatic cholangiocytes form the lining of bile ducts, augment bile composition and volume, and are characterized by expression of *SOX9*, *EPCAM*, and *KRT7*[39]. Large cholangiocytes (CH-0) are found in large bile ducts and are defined by high expression of *NCAM1*, *CFTR*, *CXCL8*, *SLC12A2*, and *CREB*. Small cholangiocytes (CH-1) are characterized by reduced expression of *CFTR* and *NCAM1*, as well as high expression of *CASR*, *GABRP*, *SLC17A4*, *SLC2A2*, and *PRSS12*[40,41].

### Supplementary Figures

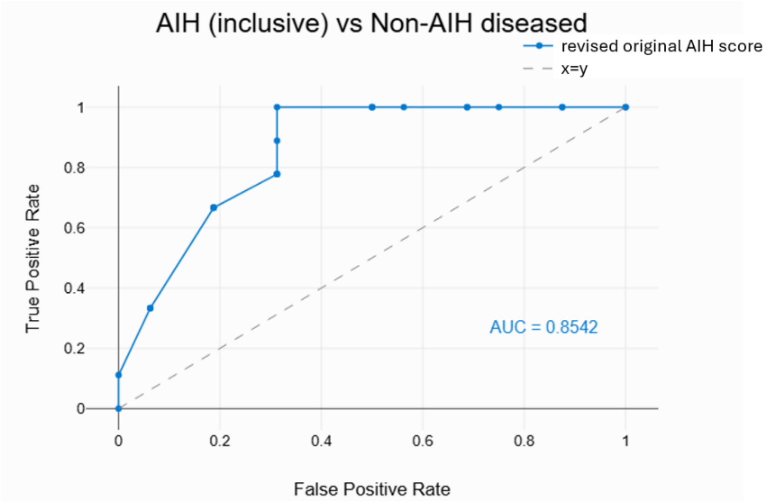


**Supplementary Figure 1:** Test characteristics of the revised original AIH score in this prospective study of possible AIH at time of liver biopsy.

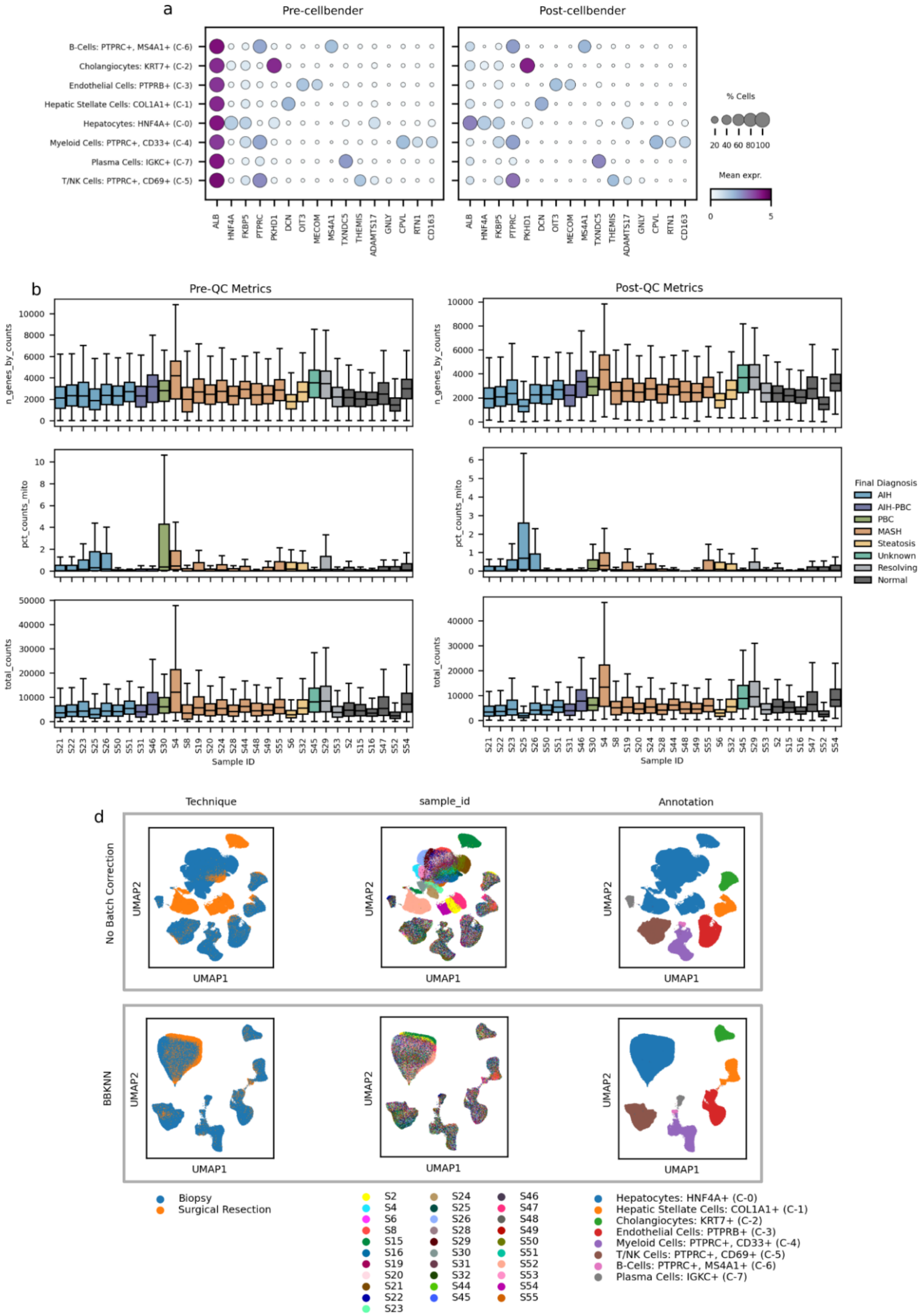

a
Pre-cellbender
Post-cellbender
B-Cells: PTPRC+, MS4A1+ (C-6)
Cholangiocytes: KRT7+ (C-2)
Endothelial Cells: PTPRB+ (C-3)
Hepatic Stellate Cells: COL1A1+ (C-1)
Hepatocytes: HNF4A+ (C-0)
Myeloid Cells: PTPRC+, CD33+ (C-4)
Plasma Cells: IGKC+ (C-7)
T/NK Cells: PTPRC+, CD69+ (C-5)
ALB HNF4A FKBP5 PTPRC PKHD1 DCN OIT3 MECOM MS4A1 TXNDC5 THEMIS ADAMTS17 GNLY CPVL RTN1 CD163
% Cells
20 40 60 80 100
Mean expr.
0 5
b
Pre-QC Metrics
Post-QC Metrics
n_genes_by_counts
pct_counts_mito
total_counts
Sample ID
S21 S22 S23 S25 S26 S50 S51 S31 S46 S30 S4 S8 S19 S20 S24 S28 S44 S48 S49 S55 S6 S32 S45 S29 S53 S2 S15 S16 S47 S52 S54
Final Diagnosis
AIH
AIH-PBC
PBC
MASH
Steatosis
Unknown
Resolving
Normal
d
Technique
sample_id
Annotation
No Batch Correction
BBKNN
UMAP1
UMAP2
Biopsy
Surgical Resection
S2 S4 S6 S8 S15 S16 S19 S20 S21 S22 S23 S24 S25 S26 S28 S29 S30 S31 S32 S44 S45 S46 S47 S48 S49 S50 S51 S52 S53 S54 S55
Hepatocytes: HNF4A+ (C-0)
Hepatic Stellate Cells: COL1A1+ (C-1)
Cholangiocytes: KRT7+ (C-2)
Endothelial Cells: PTPRB+ (C-3)
Myeloid Cells: PTPRC+, CD33+ (C-4)
T/NK Cells: PTPRC+, CD69+ (C-5)
B-Cells: PTPRC+, MS4A1+ (C-6)
Plasma Cells: IGKC+ (C-7)

**Supplementary Figure 2: Single nucleus data pre-processing.** Dotplots of select gene expression across coarse cell type clusters (a) before (left) and after (right) running ambient removal. Boxplots of quality control metrics (number of genes (top), percent of counts from mitochondrial genes (middle), and total number of counts (bottom)) per nucleus grouped by sample (b) before (left) and after (right) gene and nuclei filtering. UMAPs of all nuclei (d) without batch correction (top) and after batch-balanced-k-nearest-neighbors (BBKNN) (bottom).

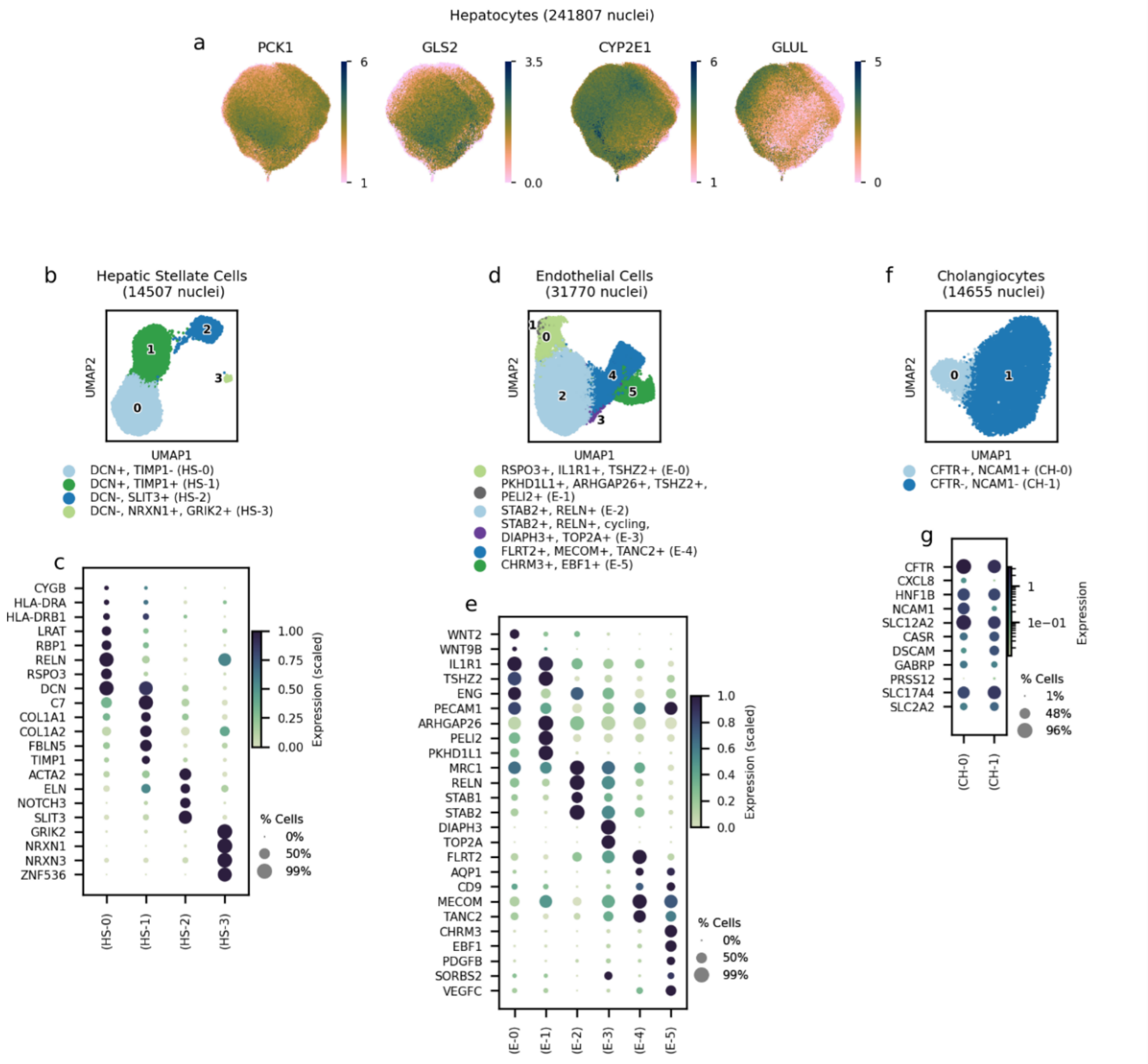


**Supplementary Figure 3: Annotation of parenchymal and non-immune cell types.** Hepatocyte UMAPs for select zonal marker genes (a). UMAPs for hepatic stellate (b), endothelial (d), and cholangiocyte (f) lineages colored by fine-grained subtypes. Marker gene dotplot for hepatic stellate cells (b), endothelial cells (e), and cholangiocytes (h). Gene expression is min-max normalized across samples and goes from green-blue (low expression) to dark blue (high expression).

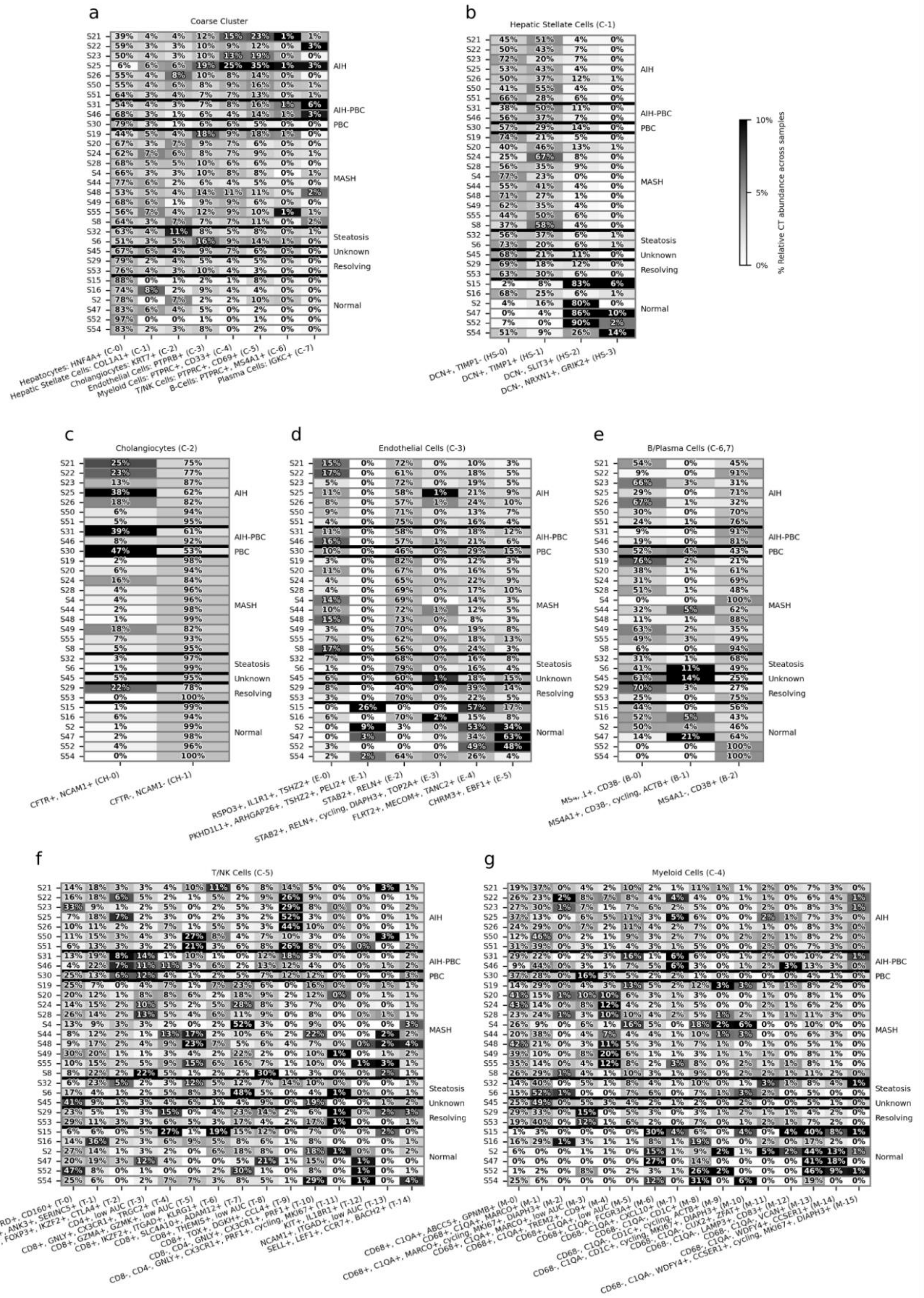

a
Coarse Cluster
Hepatocytes: HNF4A+ (C-0)
Hepatic Stellate Cells: COL1A1+ (C-1)
Cholangiocytes: KRT7+ (C-2)
Endothelial Cells: PTPRB+ (C-3)
Myeloid Cells: PTPRC+, CD33+ (C-4)
T/NK Cells: PTPRC+, CD69+ (C-5)
B-Cells: PTPRC+, MS4A1+ (C-6)
Plasma Cells: IGKC+ (C-7)
b
Hepatic Stellate Cells (C-1)
DCN+, TIMP1- (HS-0)
DCN+, TIMP1+ (HS-1)
DCN-, SLIT3+ (HS-2)
DCN-, NRXN1+, GRIK2+ (HS-3)
% Relative CT abundance across samples
10%
5%
0%
c
Cholangiocytes (C-2)
CFTR+, NCAM1+ (CH-0)
CFTR-, NCAM1- (CH-1)
d
Endothelial Cells (C-3)
RSPO3+, IL1R1+, TSHZ2+ (E-0)
PKHD1L1+, ARHGAP26+, TSHZ2+, PELI2+ (E-1)
STAB2+, RELN+ (E-2)
STAB2+, RELN+, cycling, DIAPH3+, TOP2A+ (E-3)
FLRT2+, MECOM+, TANC2+ (E-4)
CHRM3+, EBF1+ (E-5)
e
B/Plasma Cells (C-6,7)
MS4A1+, CD38- (B-0)
MS4A1+, CD38-, cycling, ACTB+ (B-1)
MS4A1-, CD38+ (B-2)
f
T/NK Cells (C-5)
g
Myeloid Cells (C-4)
AIH
AIH-PBC
PBC
MASH
Steatosis
Unknown
Resolving
Normal

**Supplementary Figure 4: Cell type abundances were calculated for across coarse cell types and subtypes.** Heatmaps for coarse cell types (a), hepatic stellate subtypes (b), cholangiocyte subtypes (c), endothelial subtypes (d), B/plasma subtypes (e), T/NK subtypes (f), and myeloid subtypes (g). Sample IDs are labeled on the left y-axis with corresponding Final Diagnosis on the right y-axis. Cell types are labelled on the x-axis. Heatmap annotated values represent cell type percent abundance relative to the number of cells in the major cell type (labelled in the title) in each sample. Coarse cell type abundances are relative to total number of cell in each sample. Heatmap hue represents a cell type abundance normalized by the sum of abundances in a cell type. This helps identify which samples have a higher abundance of a cell type relative to other samples regardless of a cell type's abundance relative to other cell types. Cell types with a light hue are less represented in that sample relative to other samples. Cell types with a dark hue are more represented in that sample.

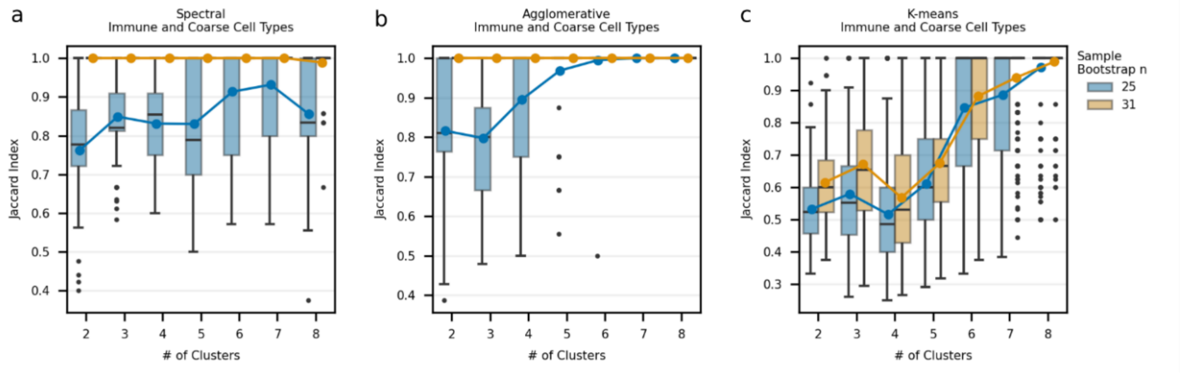


**Supplementary Figure 5: Jaccard index informs clustering consistency with varying number of clusters.** Boxplots of pairwise Jaccard index scores for spectral (n=100 runs per box) (a), agglomerative (n=100) (b) and k-means (n=500) (c) clustering of coarse and immune CTAs. Blue box plots and trend line show subsampling of n=25 samples per run with varying random seeds. Orange box plots and trend line show no subsampling of samples with varying random seeds. Outliers are shown as black dots. Blue and orange dots show the mean value for the corresponding boxplot.

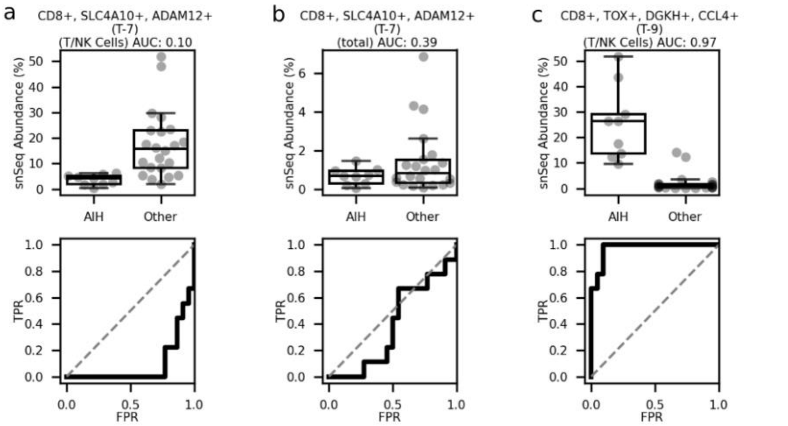


**Supplementary Figure 6: AIH samples show depletion of MAIT cells.** (a-c, top row) AIH versus non-AIH (other) fractional abundance of specific cell types divided by: total cells (b) or T/NK cells (a,c) (per sample); using the inclusive definition of AIH. (a-c, bottom row) Receiver operating characteristic curves for each cell type, with area under the curve (AUC) in the title indicating overall performance.

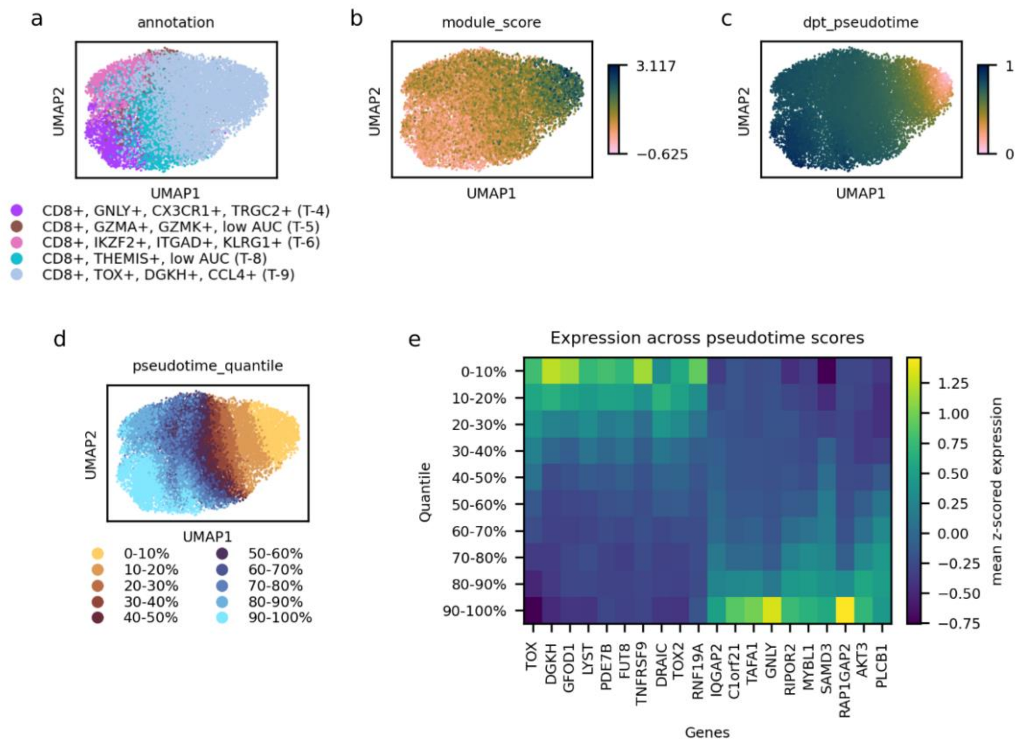


**Supplementary Figure 7: Pseudotime analysis suggests gradient of CD8+ cell phenotype.** (a-d) UMAPs of T-4,5,6,8,9 sub-clustered with T/NK annotations (a); score values from scanpy.tl.gene_score module score of genes: (TOX, CD200, TNFRSF9, CCL4) (b); scanpy.tl.dpt pseudotime values with root cell determined by cell with highest module_score (c); cells grouped by dpt_pseudotime quantiles in 10% increments (d). Top 10 genes most positively correlated (spearman correlation) and top 10 most negatively correlated are selected. Heatmap shows expression of selected genes across quantiles (e) Gene expression was z-scored across all T-4,5,6,8,9 cells and averaged (mean) within each quantile.

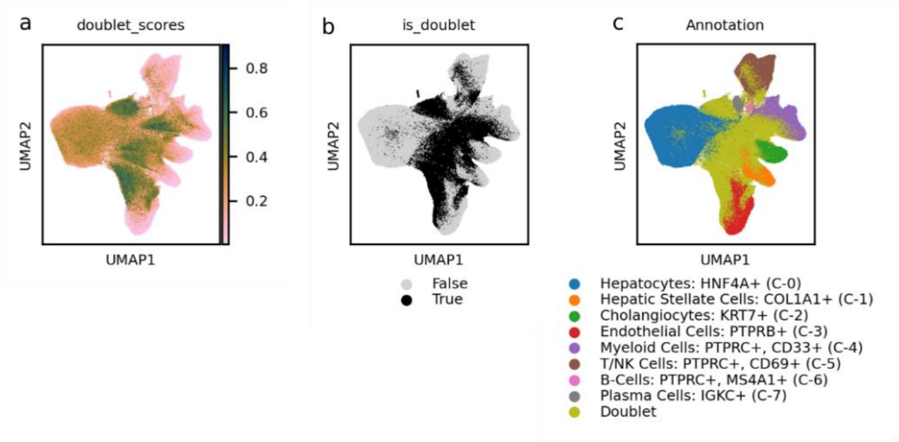


**Supplementary Figure 8: Doublet identification was supported by scrublet.** UMAPs of all nuclei including doublets (a-c) colored by scrublet assigned probabilities (a); binary label of whether a nucleus was determined a doublet (b); and coarse cell type annotations including binary doublet label (c).

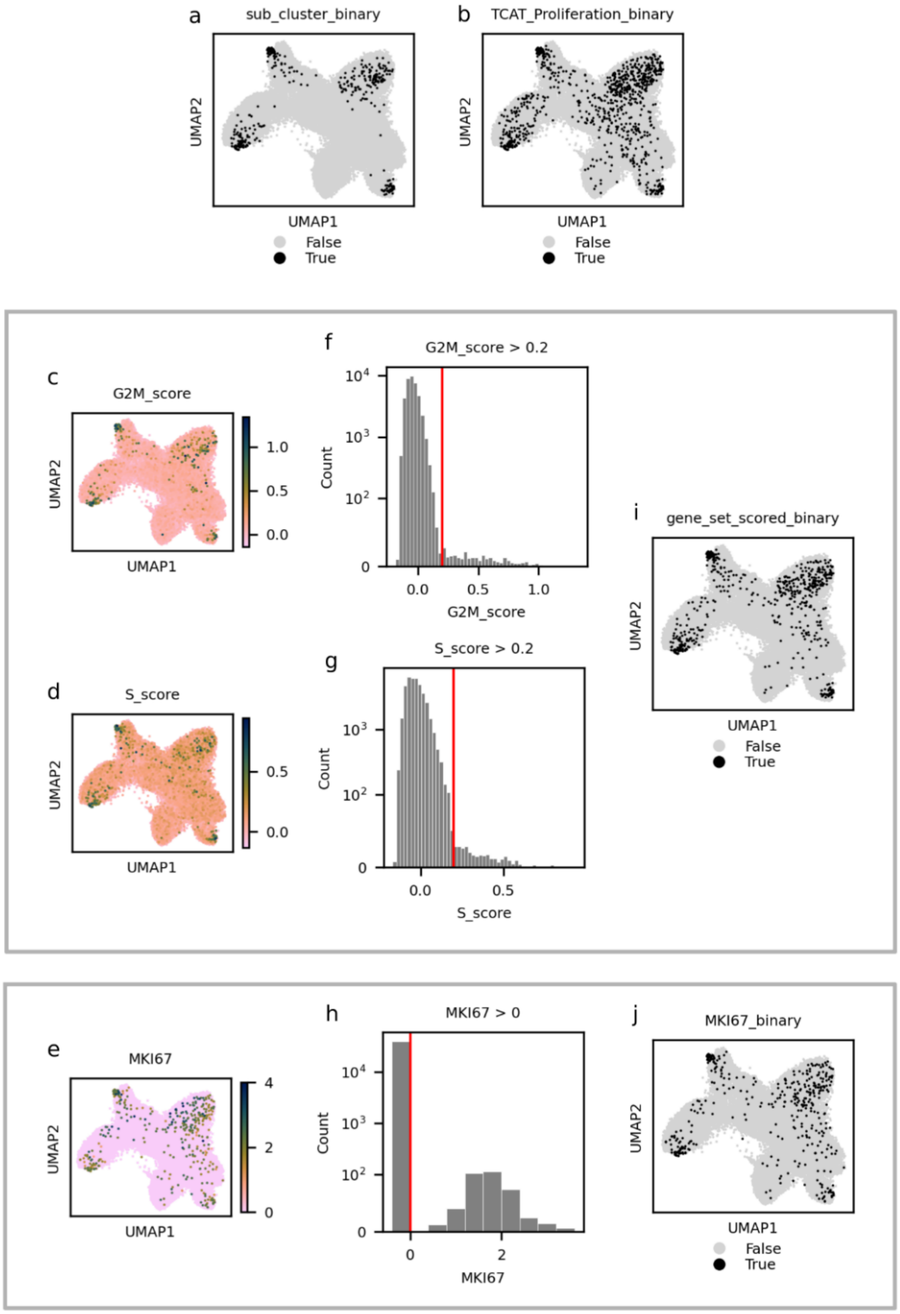


Supplementary Figure 9: **Identification of proliferating cells in snSeq data.** UMAPs of identified proliferating cells with sub-clustering (a); TCAT (b); scanpy.tl.score_genes_cell_cycle with gene sets from Tirosh *et al.* (i) and MKI67 expression (j). Black colored nuclei represent cells predicted as proliferating. UMAP of module scores of G2M phase gene set (c) and S phase gene set (d). Histogram of G2M phase gene set module scores (f), S phase gene set module scores (g), and MKI67 expression (h). Red vertical line represents cutoff value. Nuclei with scores or expression to the right of the cutoff were identified as proliferating cells for the corresponding method.

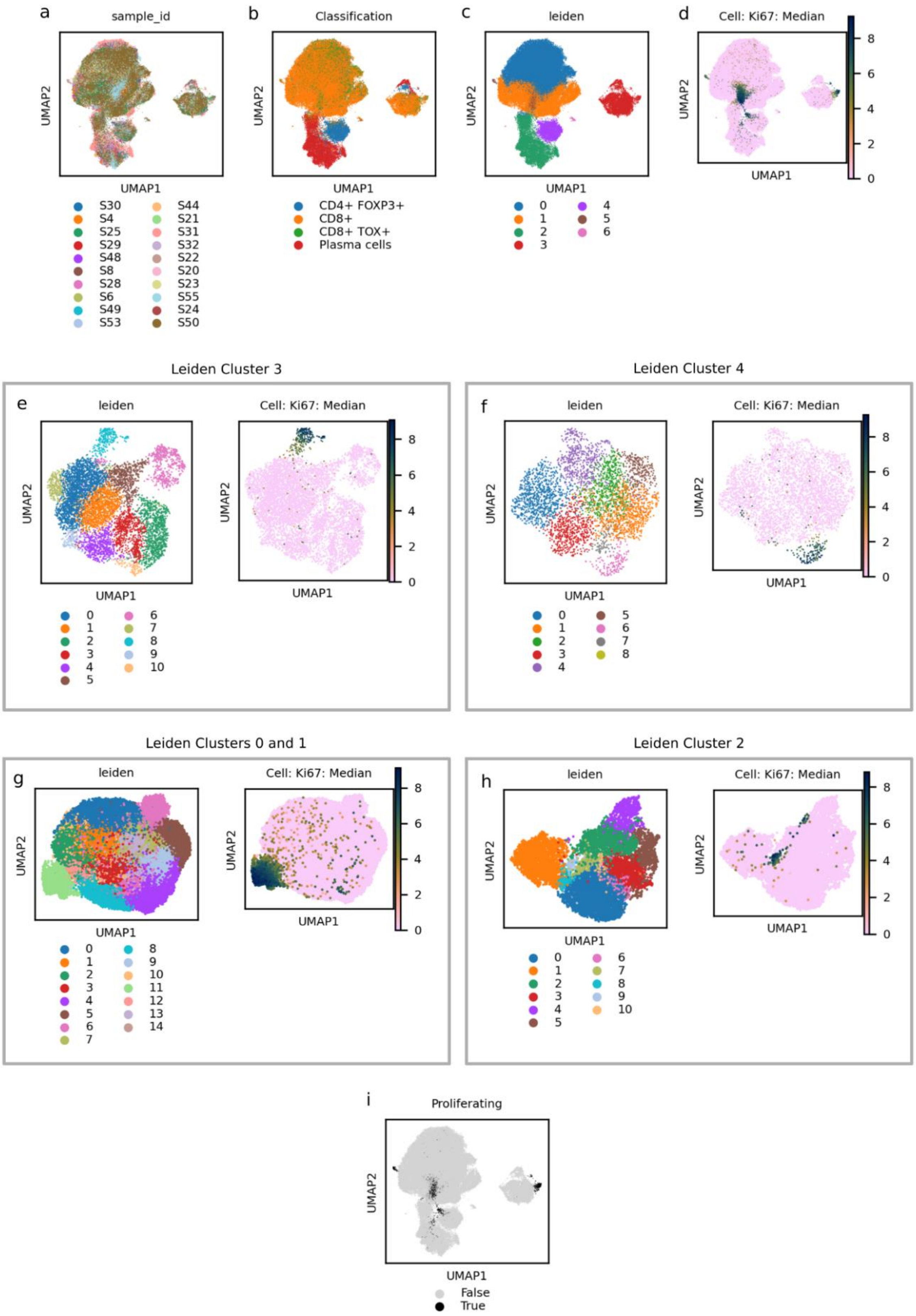

a
sample_id
UMAP1
UMAP2
S30 S4 S25 S29 S48 S8 S28 S6 S49 S53 S44 S21 S31 S32 S22 S20 S23 S55 S24 S50
b
Classification
CD4+ FOXP3+
CD8+
CD8+ TOX+
Plasma cells
c
leiden
0 1 2 3 4 5 6
d
Cell: Ki67: Median
Leiden Cluster 3
e
Leiden Cluster 4
f
Leiden Clusters 0 and 1
g
Leiden Cluster 2
h
i
Proliferating
False
True

**Supplementary Figure 10: *In situ* proliferating cell identification.** UMAPs from PhenoCycler spatial data of all classified cells (a-d,i). UMAPs of sub clustered classified cells selected by leiden cluster (e-h). Inferred proliferating cells (i). Black colored cells indicated a cell predicted as proliferating.
